## Supplemental Information for "Hippocampal-entorhinal cognitive maps and cortical motor system represent action plans and their outcomes"


| | |
|---|---|
| Supplementary Figure 1 | Goal-directed Action Task |
| Supplementary Figure 2 | Procrustes distances obtained from the two Rating Tasks |
| Supplementary Figure 3 | Behavioural performance in the two Comparison Tasks and correlation between Comparison and Rating Tasks |
| Supplementary Figure 4 | Grid-like representation of the abstract action-outcome space |
| Supplementary Figure 5 | Distance representations of the abstract action-outcome space |
| Supplementary Figure 6 | Control analyses for the representation of individual actions |
| Supplementary Figure 7 | Interaction between map-like representations in the hippocampus and individual action representations in SMA |
| Supplementary Table 1 | Significant clusters of the abstract distance-based BOLD adaptation analysis of the action combinations |
| Supplementary Table 2 | Significant clusters of the abstract distance-based BOLD adaptation analysis of the landmark outcomes of actions |
| Supplementary Table 3 | Significant clusters of the action similarity-based BOLD adaptation analysis of the action combinations |
| Supplementary Table 4 | Whole-brain clusters of the generalized Psychophysiological Interaction (gPPI) effect with left HPC used as a seed region |
| Supplementary Table 5 | Whole-brain clusters of the generalized Psychophysiological Interaction (gPPI) effect with SMA used as a seed region |

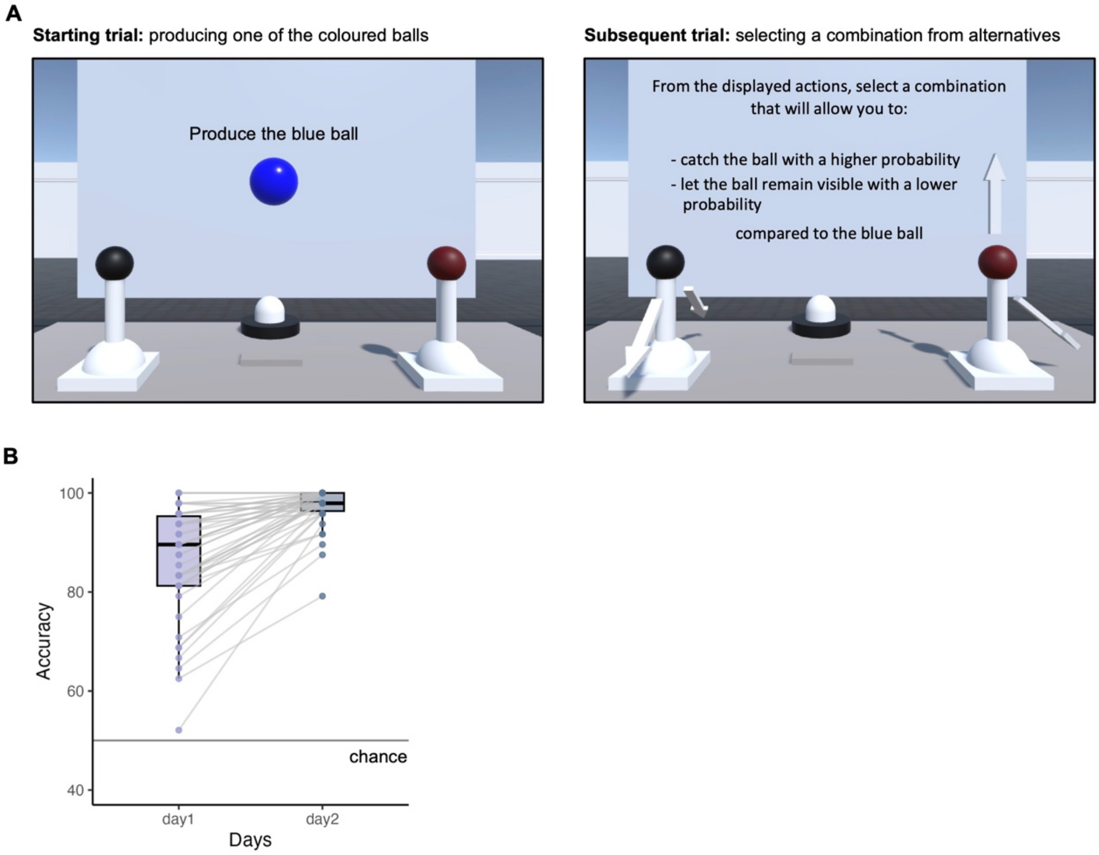


**Supplementary Figure 1: Goal-directed Action Task. A** In this VR multiple-choice task, participants completed pairs of trials. In the first, starting trial of each pair, they were instructed to produce one of the coloured balls, e.g. the blue ball. The second subsequent trial asked participants to use the previously produced coloured ball, in this case the blue ball, as a reference point and to select a correct combination of actions from the cued alternatives in order to achieve task-relevant outcomes. The example instructions are shown in the figures. **B** Behavioural performance over the two days of training, demonstrating that participants could correctly identify and perform a combination of actions from the cued alternatives to elicit the desired outcomes. Dots indicate data from individual participants.

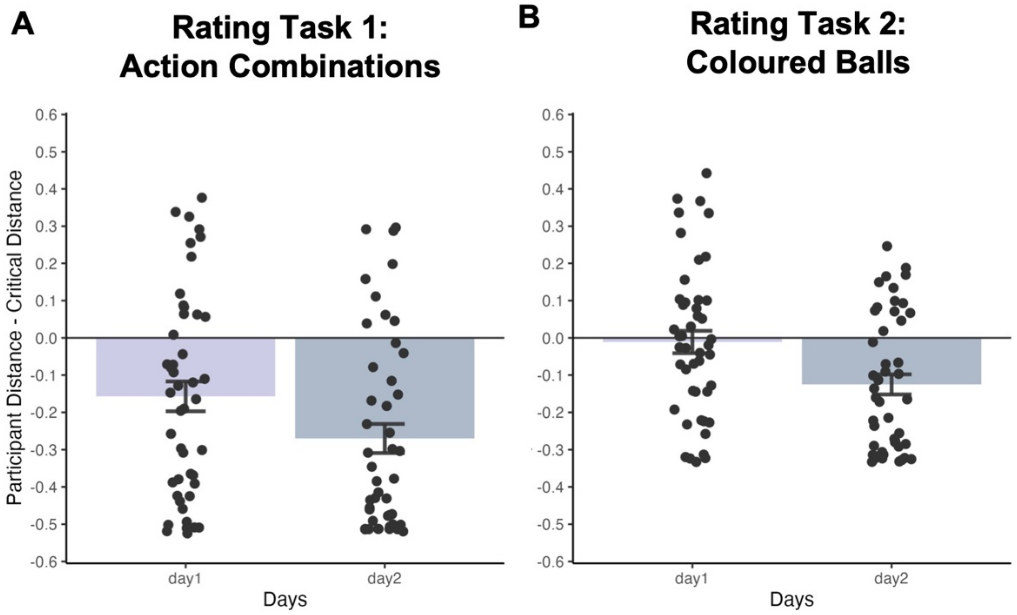


**Supplementary Figure 2: Procrustes distances obtained from the two Rating Tasks.** The Procrustes distances obtained by fitting the MDS coordinates of the action combinations (**A**) and coloured balls (**B**) to the true coordinates of the respective stimuli in the action-outcome space were smaller than the critical distances obtained from the permutation tests (see Figure 2; see Methods).

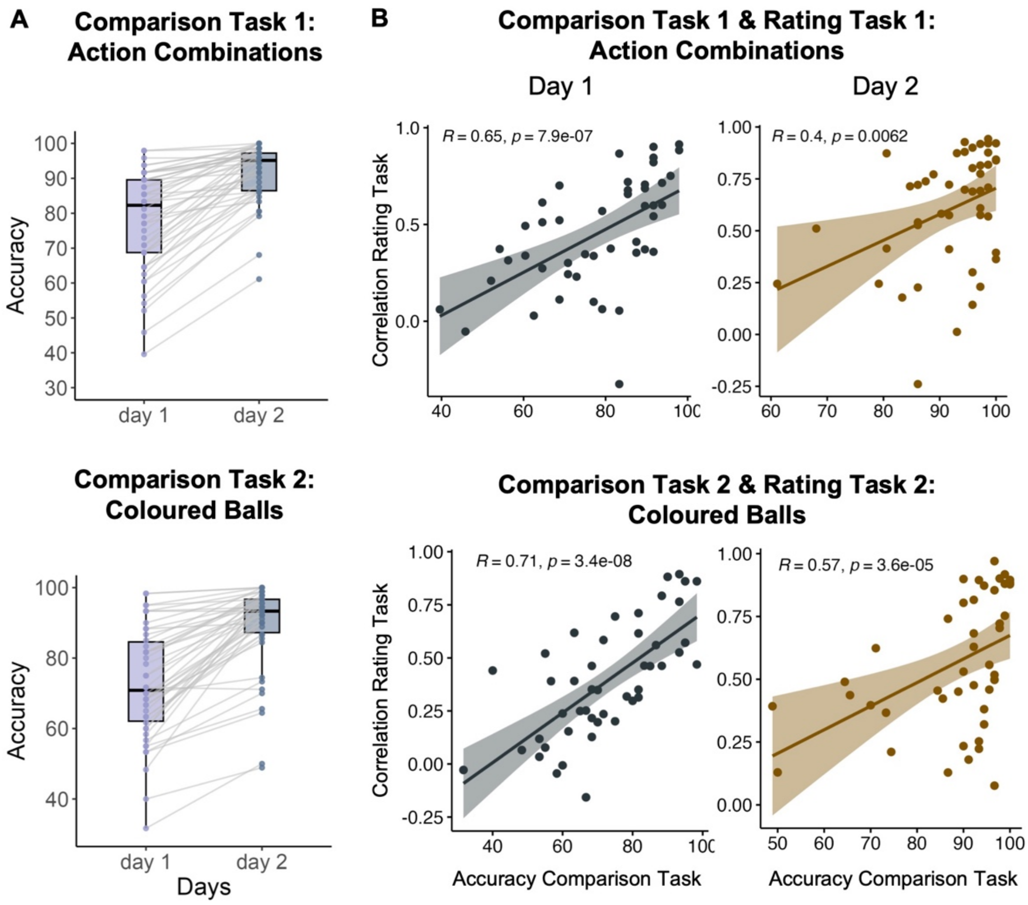


**Supplementary Figure 3: Behavioural performance in the two Comparison Tasks and correlation between Comparison and Rating Tasks. A** Overall performance over the two days of training in the Comparison tasks with action combinations (plot above) and coloured balls (plot below). **B** We examined the Spearman's correlation between performance in the Comparison and Rating Tasks with action combinations (plots above) and coloured balls (plots below) over the two training days. Accuracy in the Comparison Tasks was correlated with the correlation values obtained by matching the similarity estimates from the Rating Tasks to the action-outcome space (see Figure 2A,C and Methods).

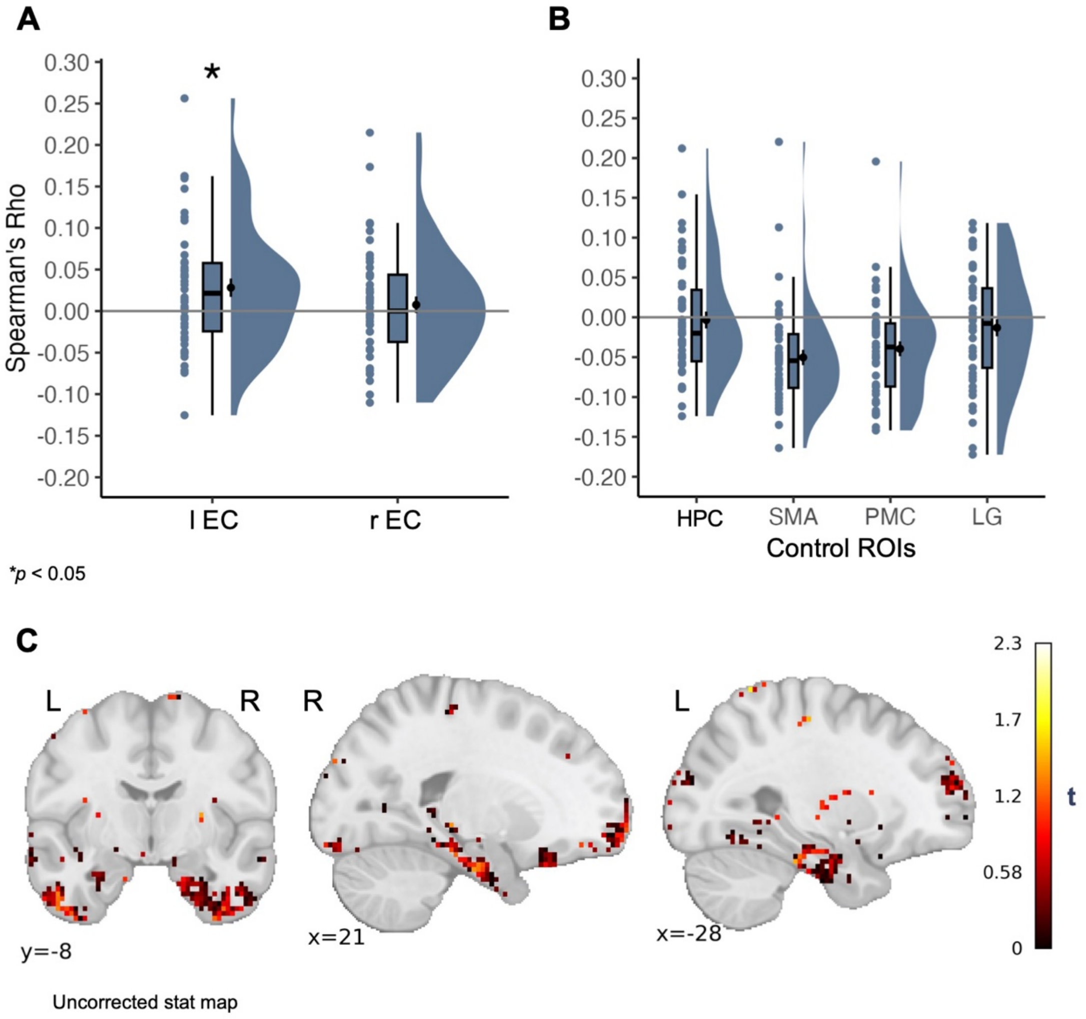


**Supplementary Figure 4: Grid-like representation of the abstract action-outcome space.** **A** Representation similarity analysis (RSA) performed on the pattern similarities from the first Comparison Task (see Methods). Here, pattern similarities were extracted separately for each hemisphere using ROI masks and showed a significant 6-fold periodicity effect in the left EC (see Results). **B** The 6-fold periodicity effect was not observed in any of the control ROIs (HPC: z(45)= -0.852, p= 0.804; SMA: z(45)= -4.638, p= 1; PMC: z(45)= -4.097, p= 1; one-tailed Wilcoxon signed rank test; LG: t(45)= -1.22, p= 0.886; one-tailed t-test). **C** Whole-brain searchlight RSA using 5 voxel radius spheres revealed a cluster in bilateral EC. Results are uncorrected for multiple comparisons. *$p < 0.05$; Bonferroni corrected for tests in both ROIs.

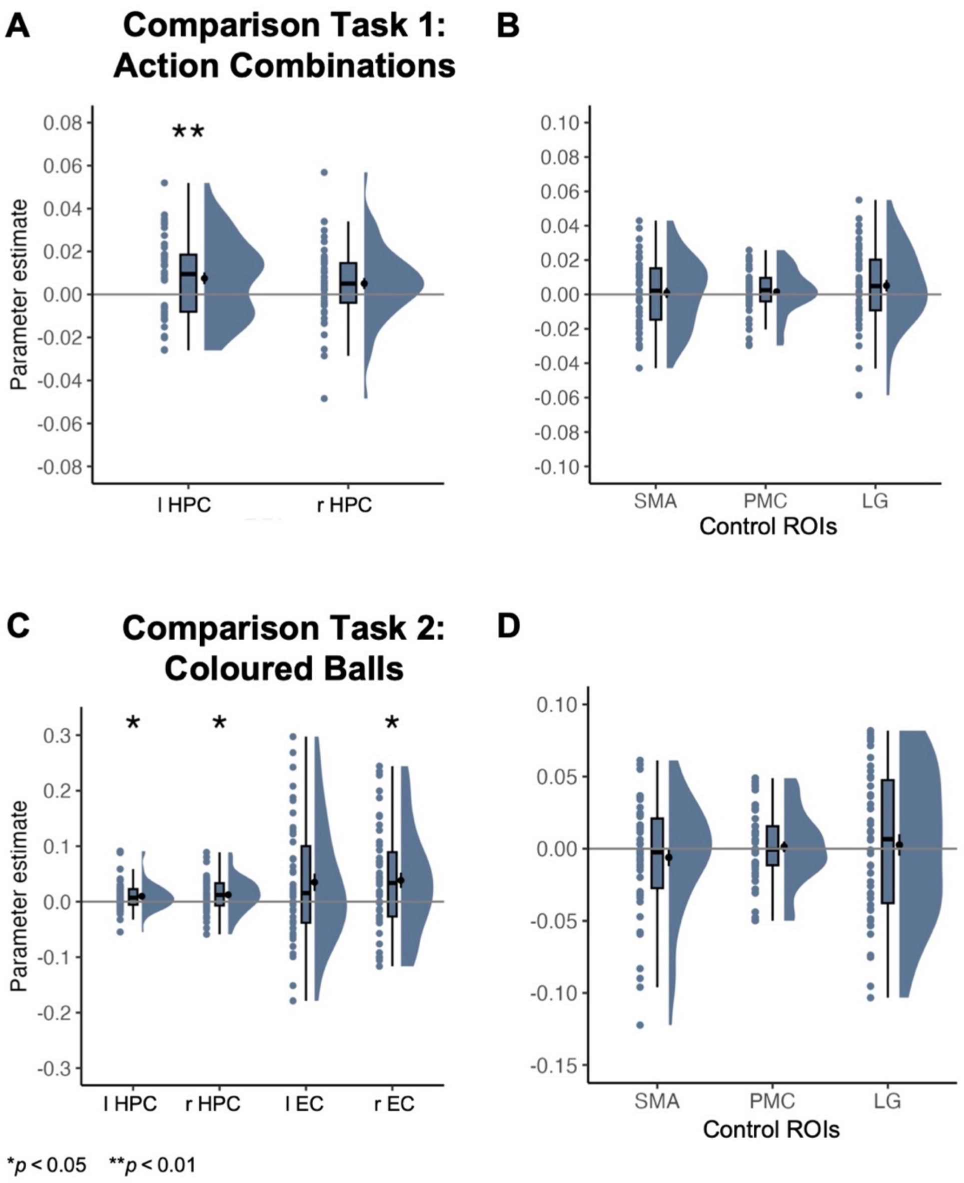


**Supplementary Figure 5: Distance representations of the abstract action-outcome space. A** The hippocampal BOLD response showed adaptation for trials with shorter distances between action combinations in the left hippocampus for the first Comparison Task (similar to the grid-like representation, see Supplementary Figure 4A; see Results). **B** None of the control ROIs showed any adaptation of the BOLD signal to the distance between action combinations in the abstract action-outcome space (SMA: t(45)= 0.307, p= 0.38; PMC: t(45)= 0.808, p= 0.212; LG: t(45)= 1.52, p= 0.067; one-tailed t-test). **C** The adaptation effect in the second Comparison Task, for coloured balls positioned closer to each other in the action-outcome space, was present in both hemispheres for the HPC and only in the right hemisphere for the EC (Left HPC: z(45) = 2.631, p= 0.033; one-tailed Wilcoxon signed rank test; Right HPC: t(45)= 2.70, p= 0.019; Left EC: t(45)= 2.21, p= 0.064; Right EC: t(45)= 2.74, p= 0.017; one-tailed t-test). **D** The control ROIs including SMA, PMC and LG did not show the adaptation effect (SMA: z(45)= 0.401, p= 0.656; one-tailed Wilcoxon signed rank test; PMC: t(45)= 0.347, p= 0.365; LG: t(45)= 0.352, p= 0.363; one-tailed t-test). *$p < 0.05$; **$p < 0.01$; The results for the HPC and EC ROIs are Bonferroni corrected for tests in multiple ROIs.

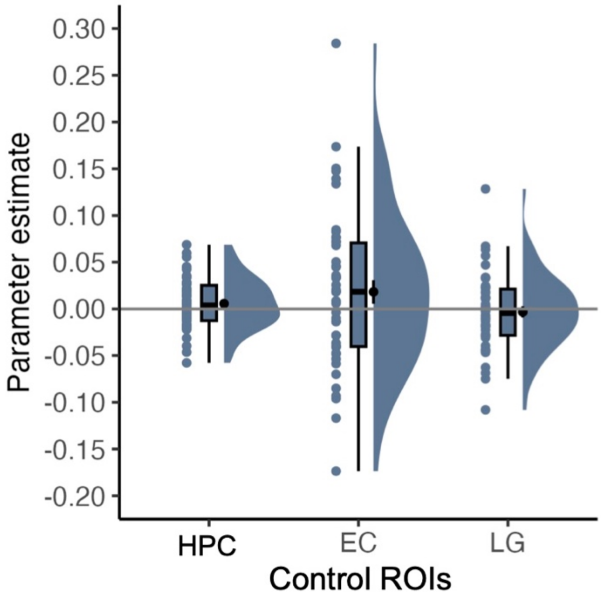


**Supplementary Figure 6: Control analyses for the representation of individual actions.** The shared actions between action combinations did not elicit the modulation of BOLD response in the control ROIs (HPC: t(45)= 1.40, p= 0.168; EC: t(45)= 1.45, p= 0.153; LG: t(45)= -0.531, p= 0.598; two-tailed t-test).

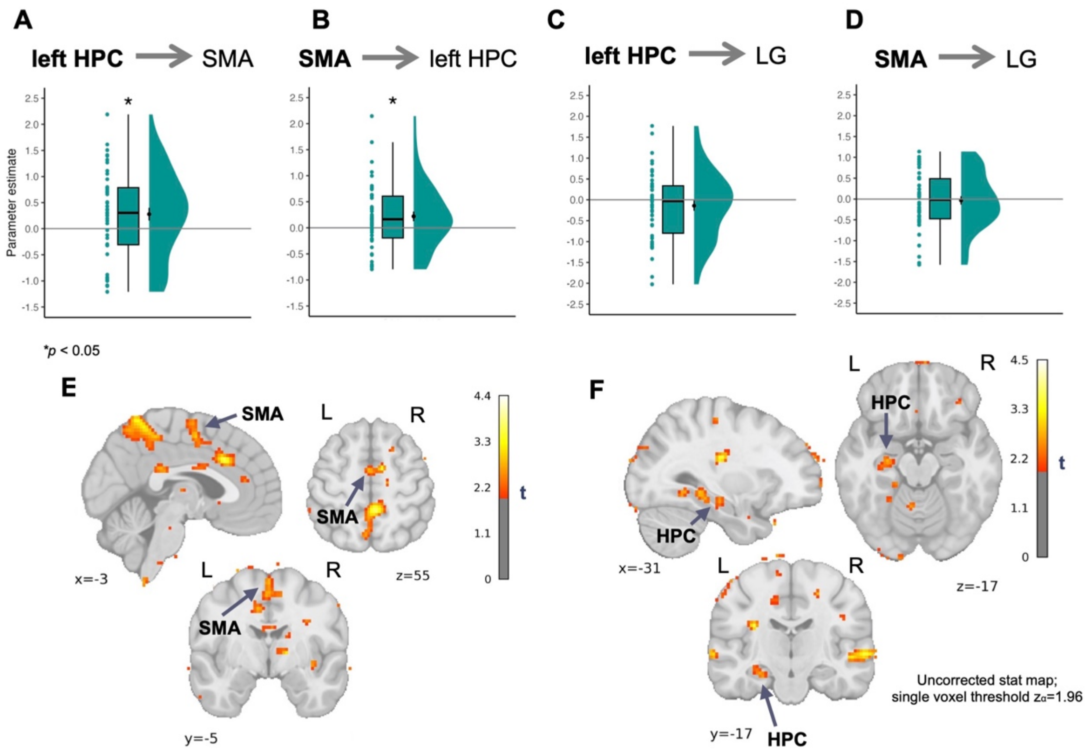


**Supplementary Figure 7: Interaction between map-like representations in the hippocampus and individual action representations in SMA.** Analyses were performed on the data from the first Comparison Task (see Methods). **A,B,C,D** left HPC used as a seed ROI to assess the connectivity with the SMA (HPC-SMA: t(45) = 2.28, p= 0.027; two-tailed t-test) (**A**) and with the control ROI LG (HPC-LG: t(45)= -1.15; p= 0.256; two-tailed t-test) (**C**). SMA used as a seed ROI to assess the connectivity with the left HPC (SMA-HPC: t(45)= 2.40; p= 0.02; two-tailed t-test) (**B**) and with the control ROI LG (SMA-LG: t(45)= -0.344; p=0.733; two-tailed t-test) (**D**). **E** Whole-brain connectivity map for the left HPC used as a seed. MNI peak voxel coordinates of SMA: 3, 1, 48; peak voxel t(45) = 3.175; two-tailed test. **F** Whole-brain connectivity map for the SMA used as a seed. MNI peak voxel coordinates of lHC: -26, -15, -18; peak voxel t(45) = 2.802; two-tailed test. Whole-brain data is uncorrected for multiple comparisons. Statistical significance threshold is defined at a single voxel level ($z_\alpha$=1.96). **p* < 0.05.

**Supplementary Table 1: Significant clusters of the abstract distance-based BOLD adaptation analysis of the action combinations.** Clusters surviving the whole-brain correction using FDR method with a voxel-level threshold of p<0.01. Table displays MNI coordinates (X, Y, Z), statistical T values and atlas labels of peak voxels of the clusters. Atlas labels are based on the Juelich Histological Atlas (JHA) and Harvard-Oxford Cortical Structural Atlas (HOCSA). The labels were generated using FSL. Atlases are listed only if the labels were found for a given atlas. The letters after the cluster ID indicates its subcluster.

| Cluster ID | X | Y | Z | T value | Cluster Size (voxels) | Atlas label |
|---|---|---|---|---|---|---|
| 1 | -36.788 | -35.468 | 12.25 | 6.668 | 1446 | JHA: |

| | | | | | | |
|---|---|---|---|---|---|---|
| | | | | | | 26% GM Primary auditory cortex TE1.1 L, 18% WM Acoustic radiation L, 8% WM Optic radiation L, 6% GM Secondary somatosensory cortex / Parietal operculum OP1 L, 6% GM Inferior parietal lobule PFcm L, 2% GM Insula Ig1 L<br><br>HOCSA:<br>34% Planum Temporale, 2% Supramarginal Gyrus, posterior division, 1% Heschl's Gyrus (includes H1 and H2), 1% Parietal Operculum Cortex |
| 1a | -44.252 | -30.492 | 20.5 | 4.473 | | JHA:<br>61% GM Secondary somatosensory cortex / Parietal operculum OP1 L, 23% GM Inferior parietal lobule PFop L, 18% GM Inferior parietal lobule PFcm L, 15% GM Primary auditory cortex TE1.1 L, 4% GM Primary auditory cortex TE1.0 L, 2% GM Insula Ig2 L<br><br>HOCSA:<br>65% Parietal Operculum Cortex, 7% Central Opercular Cortex, 2% Superior Temporal Gyrus, posterior division, 1% Planum Temporale, 1% Heschl's Gyrus (includes H1 and H2) |
| 2 | 17.947 | 49.123 | 37.0 | 5.881 | 3881 | HOCSA:<br>84% Frontal Pole |
| 2a | 25.411 | 51.611 | 23.25 | 4.820 | | HOCSA:<br>82% Frontal Pole |
| 3 | 57.755 | -28.004 | 23.25 | 5.842 | 8239 | JHA:<br>49% GM Inferior parietal lobule PFcm R, 32% GM Secondary somatosensory cortex / Parietal operculum OP1 R, 12% GM Inferior parietal lobule PF R, 4% GM Inferior parietal lobule PFt R, 1% GM Inferior parietal lobule PFop R<br><br>HOCSA:<br>33% Parietal Operculum Cortex, 21% Supramarginal Gyrus, anterior division, 13% Planum Temporale, 1% Supramarginal Gyrus, posterior division |
| 3a | 67.707 | -40.444 | 26.0 | 5.351 | | JHA:<br>22% GM Inferior parietal lobule PF R, 2% GM Inferior parietal lobule Pga R<br><br>HOCSA:<br>34% Supramarginal Gyrus, posterior division, 2% Angular Gyrus, 1% Superior Temporal Gyrus, posterior division |
| 3b | 67.707 | -47.908 | 9.5 | 5.284 | | JHA:<br>3% GM Inferior parietal lobule Pga R<br><br>HOCSA:<br>24% Middle Temporal Gyrus, temporooccipital part, 11% Angular Gyrus, 4% Supramarginal Gyrus, posterior division |
| 3c | 65.219 | -32.98 | 28.75 | 5.212 | | JHA:<br>83% GM Inferior parietal lobule PF R, 17% GM Inferior parietal lobule PFcm R<br><br>HOCSA:<br>33% Supramarginal Gyrus, anterior division, 13% Superior Temporal Gyrus, posterior division, 11% Supramarginal Gyrus, posterior division, 9% Planum Temporale, 7% Parietal Operculum Cortex |
| 4 | -24.348 | 49.123 | 37.0 | 5.640 | 731 | HOCSA:<br>77% Frontal Pole |
| 5 | 52.779 | -0.636 | 9.5 | 5.595 | 2059 | JHA:<br>25% GM Secondary somatosensory cortex / Parietal operculum OP4 R, 14% GM Secondary somatosensory cortex / Parietal operculum OP3 R, 2% GM Inferior parietal lobule PFop R<br><br>HOCSA:<br>34% Central Opercular Cortex, 9% Precentral Gyrus, 1% Inferior Frontal Gyrus, pars opercularis |

| | | | | | | |
|---|---|---|---|---|---|---|
| 5a | 45.315 | 6.827 | 6.75 | 4.34 | | JHA:<br>6% GM Secondary somatosensory cortex / Parietal operculum OP4 R, 1% GM Broca's area BA44 R<br><br>HOCSA:<br>54% Central Opercular Cortex, 5% Frontal Operculum Cortex, 2% Precentral Gyrus, 1% Inferior Frontal Gyrus, pars opercularis |
| 6 | 50.291 | -18.052 | -12.5 | 5.587 | 1038 | JHA:<br>1% GM Insula Id1 R<br><br>HOCSA:<br>51% Middle Temporal Gyrus, posterior division, 20% Superior Temporal Gyrus, posterior division, 2% Middle Temporal Gyrus, anterior division |
| 6a | 57.755 | -13.076 | -7.0 | 4.3 | | HOCSA:<br>43% Superior Temporal Gyrus, posterior division, 28% Middle Temporal Gyrus, posterior division, 7% Middle Temporal Gyrus, anterior division, 3% Superior Temporal Gyrus, anterior division |
| 6b | 65.219 | -18.052 | -7.0 | 3.882 | | HOCSA:<br>59% Middle Temporal Gyrus, posterior division, 19% Superior Temporal Gyrus, posterior division, 2% Middle Temporal Gyrus, anterior division |
| 7 | -39.276 | 1.851 | 15.0 | 5.542 | 800 | JHA:<br>6% GM Broca's area BA44 L<br><br>HOCSA:<br>30% Central Opercular Cortex |
| 8 | -1.956 | -18.052 | 42.5 | 5.147 | 2893 | JHA:<br>9% GM Primary motor cortex BA4a L, 6% GM Premotor cortex BA6 L<br><br>HOCSA:<br>65% Cingulate Gyrus, posterior division, 20% Cingulate Gyrus, anterior division, 7% Precentral Gyrus |
| 8a | 0.531 | -3.124 | 45.25 | 4.785 | | JHA:<br>36% GM Premotor cortex BA6 L<br><br>HOCSA:<br>43% Cingulate Gyrus, anterior division, 35% Juxtapositional Lobule Cortex (formerly Supplementary Motor Cortex), 1% Cingulate Gyrus, posterior division |
| 8b | 7.995 | -20.54 | 42.5 | 4.21 | | JHA: 26% GM Premotor cortex BA6 R<br><br>HOCSA: 61% Cingulate Gyrus, posterior division, 9% Precentral Gyrus, 7% Cingulate Gyrus, anterior division |
| 8c | -11.908 | -10.588 | 50.75 | 4.017 | | JHA: 43% GM Premotor cortex BA6 L<br><br>HOCSA: 17% Juxtapositional Lobule Cortex (formerly Supplementary Motor Cortex), 1% Precentral Gyrus |
| 9 | -9.42 | -90.204 | 34.25 | 5.022 | 527 | JHA: 9% GM Superior parietal lobule 7P L, 2% GM Visual cortex V1 BA17 L, 1% GM Visual cortex V2 BA18 L, 1% GM Superior parietal lobule 7A L<br><br>HOCSA: 44% Occipital Pole, 19% Lateral Occipital Cortex, superior division, 5% Cuneal Cortex |
| 10 | 50.291 | 39.171 | -1.5 | 4.894 | 1055 | JHA: 11% GM Broca's area BA45 R<br><br>HOCSA: 73% Frontal Pole, 10% Inferior Frontal Gyrus, pars triangularis |
| 10a | 57.755 | 36.683 | 1.25 | 4.555 | | JHA: 39% GM Broca's area BA45 R<br><br>HOCSA: 14% Frontal Pole, 6% Inferior Frontal Gyrus, pars triangularis |

| | | | | | | |
|---|---|---|---|---|---|---|
| 11 | 30.387 | -23.028 | -20.75 | 4.87 | 425 | JHA:<br>37% GM Hippocampus subiculum R, 21% GM Hippocampus cornu ammonis R, 7% WM Cingulum R, 2% WM Optic radiation R, 2% GM Hippocampus dentate gyrus R<br><br>HOCSA: 32% Parahippocampal Gyrus, posterior division, 6% Parahippocampal Gyrus, anterior division, 1% Temporal Fusiform Cortex, posterior division |
| 12 | -11.908 | -37.956 | 42.5 | 4.84 | 459 | JHA:<br>17% GM Superior parietal lobule 5Ci L, 5% GM Superior parietal lobule 5M L<br><br>HOCSA:<br>29% Precuneous Cortex, 29% Cingulate Gyrus, posterior division, 6% Precentral Gyrus, 5% Postcentral Gyrus |
| 13 | -64.156 | -23.028 | 17.75 | 4.819 | 1498 | JHA:<br>49% GM Secondary somatosensory cortex / Parietal operculum OP1 L, 38% GM Inferior parietal lobule PFop L, 1% GM Inferior parietal lobule PFt L<br><br>HOCSA:<br>25% Supramarginal Gyrus, anterior division, 24% Postcentral Gyrus, 12% Parietal Operculum Cortex, 9% Planum Temporale, 8% Central Opercular Cortex, 3% Superior Temporal Gyrus, posterior division |
| 14 | 50.291 | 6.827 | -40.0 | 4.733 | 1293 | HOCSA:<br>57% Temporal Pole, 17% Inferior Temporal Gyrus, anterior division, 7% Middle Temporal Gyrus, anterior division |
| 15 | 60.243 | -13.076 | 15.0 | 4.688 | 442 | JHA:<br>57% GM Secondary somatosensory cortex / Parietal operculum OP1 R, 42% GM Secondary somatosensory cortex / Parietal operculum OP4 R, 22% GM Inferior parietal lobule PFop R, 12% GM Secondary somatosensory cortex / Parietal operculum OP3 R<br><br>HOCSA:<br>46% Central Opercular Cortex, 16% Postcentral Gyrus, 7% Parietal Operculum Cortex, 2% Planum Temporale, 2% Planum Polare, 1% Heschl's Gyrus (includes H1 and H2), 1% Supramarginal Gyrus, anterior division, 1% Precentral Gyrus |
| 16 | -31.812 | -15.564 | 6.75 | 4.603 | 459 | JHA:<br>4% GM Insula Ig2 L, 2% GM Secondary somatosensory cortex / Parietal operculum OP3 L<br><br>HOCSA:<br>3% Insular Cortex |
| 17 | 10.483 | -85.228 | 37.0 | 4.505 | 408 | JHA:<br>1% GM Visual cortex V2 BA18 R<br><br>HOCSA:<br>29% Cuneal Cortex, 25% Occipital Pole, 9% Lateral Occipital Cortex, superior division, 1% Precuneous Cortex |
| 18 | 30.387 | -40.444 | 67.25 | 4.457 | 323 | JHA:<br>56% GM Primary somatosensory cortex BA1 R, 34% GM Primary somatosensory cortex BA3b R, 21% GM Primary somatosensory cortex BA2 R, 10% GM Superior parietal lobule 7PC R, 10% GM Primary motor cortex BA4p R, 4% GM Premotor cortex BA6 R, 4% GM Primary motor cortex BA4a R, 3% WM Corticospinal tract R, 1% GM Superior parietal lobule 5M R, 1% GM Superior parietal lobule 5L R |

| | | | | | | |
|---|---|---|---|---|---|---|
| | | | | | | HOCSA:<br>38% Superior Parietal Lobule, 27% Postcentral Gyrus |
| 18a | 27.899 | -37.956 | 59.0 | 3.851 | | JHA:<br>52% GM Primary somatosensory cortex BA2 R, 45% GM Primary somatosensory cortex BA3b R, 10% GM Primary somatosensory cortex BA1 R, 10% GM Primary motor cortex BA4p R, 7% GM Superior parietal lobule 7A R, 5% GM Superior parietal lobule 7PC R, 3% GM Superior parietal lobule 5L R, 1% GM Superior parietal lobule 5M R<br><br>HOCSA:<br>45% Postcentral Gyrus, 22% Superior Parietal Lobule |
| 19 | 37.851 | 29.219 | -18.0 | 4.445 | 170 | HOCSA:<br>69% Frontal Orbital Cortex, 23% Frontal Pole |
| 20 | 22.923 | -55.372 | -12.5 | 4.366 | 425 | JHA:<br>4% GM Visual cortex V3V R, 1% GM Visual cortex V4 R, 1% GM Visual cortex V2 BA18 R<br><br>HOCSA:<br>48% Lingual Gyrus, 36% Temporal Occipital Fusiform Cortex, 4% Occipital Fusiform Gyrus |
| 21 | -1.956 | -13.076 | 70.0 | 4.105 | 323 | JHA:<br>76% GM Premotor cortex BA6 L, 16% GM Primary motor cortex BA4a L<br><br>HOCSA:<br>32% Precentral Gyrus, 26% Juxtapositional Lobule Cortex(formerly Supplementary Motor Cortex) |
| 21a | -4.444 | -10.588 | 61.75 | 3.912 | | JHA:<br>94% GM Premotor cortex BA6 L, 4% GM Primary motor cortex BA4a L, 2% WM Corticospinal tract L<br><br>HOCSA:<br>63% Juxtapositional Lobule Cortex (formerly Supplementary Motor Cortex), 10% Precentral Gyrus |
| 22 | -46.74 | -10.588 | 20.5 | 4.046 | 170 | JHA:<br>38% GM Secondary somatosensory cortex / Parietal operculum OP3 L, 31% GM Secondary somatosensory cortex / Parietal operculum OP4 L, 8% GM Secondary somatosensory cortex / Parietal operculum OP1 L, 3% GM Primary somatosensory cortex BA3a L<br><br>HOCSA: 13% Central Opercular Cortex |

**Supplementary Table 2: Significant clusters of the abstract distance-based BOLD adaptation analysis of the landmark outcomes of actions.** Clusters surviving the whole-brain correction using FDR method with a voxel-level threshold of p<0.01. Table displays MNI coordinates (X, Y, Z), statistical T values and atlas labels of peak voxels of the clusters. Atlas labels are based on the Juelich Histological Atlas (JHA) and Harvard-Oxford Cortical Structural Atlas (HOCSA). The labels were generated using FSL. Atlases are listed only if the labels were found for a given atlas. The letters after the cluster ID indicates its subcluster.

| Cluster ID | X | Y | Z | T Value | Cluster Size (voxels) | Atlas label |
|---|---|---|---|---|---|---|
| 1 | 7.995 | 59.075 | 9.5 | 6.896 | 19610 | HOCSA: |

| | | | | | | |
|---|---|---|---|---|---|---|
| | | | | | | 56% Frontal Pole, 3% Paracingulate Gyrus, 1% Superior Frontal Gyrus |
| 1a | 17.947 | 44.147 | 48.0 | 6.434 | | HOCSA:<br>66% Frontal Pole, 5% Superior Frontal Gyrus |
| 1b | 17.947 | 51.611 | 37.0 | 6.265 | | HOCSA:<br>83% Frontal Pole |
| 1c | 17.947 | 39.171 | 50.75 | 6.084 | | JHA:<br>6% GM Premotor cortex BA6 R<br><br>HOCSA:<br>53% Frontal Pole, 24% Superior Frontal Gyrus |
| 2 | 32.875 | 6.827 | 9.5 | 5.983 | 16835 | HOCSA:<br>28% Insular Cortex, 1% Central Opercular Cortex |
| 2a | 52.779 | 26.731 | 4.0 | 5.429 | | JHA:<br>50% GM Broca's area BA45 R, 1% GM Broca's area BA44 R<br><br>HOCSA:<br>38% Inferior Frontal Gyrus, pars triangularis, 8% Inferior Frontal Gyrus, pars opercularis, 5% Frontal Operculum Cortex, 2% Frontal Orbital Cortex |
| 2b | 45.315 | 19.267 | -12.5 | 5.365 | | HOCSA:<br>36% Frontal Orbital Cortex, 9% Temporal Pole, 3% Insular Cortex, 1% Frontal Operculum Cortex |
| 2c | 35.363 | -5.612 | 4.0 | 5.344 | | JHA:<br>8% GM Secondary somatosensory cortex / Parietal operculum OP3 R, 8% GM Secondary somatosensory cortex / Parietal operculum OP2 R<br><br>HOCSA:<br>27% Insular Cortex |
| 3 | -41.764 | 9.315 | -4.25 | 5.86 | 8170 | HOCSA:<br>53% Insular Cortex, 10% Central Opercular Cortex, 5% Frontal Operculum Cortex |
| 3a | -36.788 | -13.076 | 4.0 | 5.396 | | JHA:<br>12% GM Secondary somatosensory cortex / Parietal operculum OP3 L, 5% GM Insula Ig2 L, 3% GM Insula Id1 L<br><br>HOCSA:<br>52% Insular Cortex |
| 3b | -39.276 | -23.028 | 1.25 | 5.291 | | JHA:<br>45% WM Acoustic radiation L, 37% GM Insula Id1 L, 35% GM Insula Ig2 L, 25% GM Primary auditory cortex TE1.1 L, 10% GM Primary auditory cortex TE1.0 L, 6% GM Insula Ig1 L<br><br>HOCSA:<br>36% Heschl's Gyrus (includes H1 and H2), 20% Planum Polare, 2% Insular Cortex |
| 3c | -36.788 | -3.124 | -1.5 | 5.129 | | JHA:<br>1% WM Inferior occipito-frontal fascicle L<br><br>HOCSA:<br>24% Insular Cortex |
| 4 | 57.755 | -35.468 | 34.25 | 5.533 | 6587 | JHA:<br>63% GM Inferior parietal lobule PF R, 20% GM Inferior parietal lobule PFcm R, 16% GM Inferior parietal lobule PFm R, 12% GM Anterior intra-parietal sulcus hIP2 R<br>HOCSA:<br>30% Supramarginal Gyrus, posterior division, 14% Parietal Operculum Cortex, 11% Supramarginal Gyrus, anterior division, 4% Planum Temporale, 2% Angular Gyrus |
| 4a | 60.243 | -45.42 | 31.5 | 5.224 | | JHA:<br>78% GM Inferior parietal lobule PFm R, 20% GM Inferior parietal lobule Pga R, 16% GM Inferior parietal lobule PF R<br>HOCSA:<br>45% Angular Gyrus, 43% Supramarginal Gyrus, posterior division |
| 4b | 52.779 | -57.86 | 39.75 | 4.962 | | JHA: |

| | | | | | | |
|---|---|---|---|---|---|---|
| | | | | | | 73% GM Inferior parietal lobule Pga R, 30% GM Inferior parietal lobule PGp R, 6% GM Inferior parietal lobule PFm R, 1% GM Anterior intra-parietal sulcus hIP1 R<br>HOCSA:<br>41% Angular Gyrus, 36% Lateral Occipital Cortex, superior division |
| 4c | 60.243 | -55.372 | 42.5 | 4.873 | | JHA:<br>4% GM Inferior parietal lobule Pga R, 3% GM Inferior parietal lobule PFm R, 1% GM Inferior parietal lobule PGp R<br>HOCSA:<br>20% Angular Gyrus, 9% Lateral Occipital Cortex, superior division |
| 5 | -24.348 | -5.612 | -20.75 | 5.348 | 1702 | JHA:<br>88% GM Amygdala_laterobasal group L, 28% GM Hippocampus cornu ammonis L, 25% GM Amygdala_superficial group L, 8% GM Amygdala_centromedial group L, 4% GM Hippocampus subiculum L |
| 6 | 7.995 | -20.54 | 42.5 | 5.292 | 6758 | JHA:<br>26% GM Premotor cortex BA6 R<br>HOCSA:<br>61% Cingulate Gyrus, posterior division, 9% Precentral Gyrus, 7% Cingulate Gyrus, anterior division |
| 6a | 5.507 | -32.98 | 50.75 | 4.856 | | JHA:<br>48% GM Primary motor cortex BA4a R, 42% GM Superior parietal lobule 5M R, 24% GM Superior parietal lobule 5Ci R, 7% WM Corticospinal tract R, 4% GM Premotor cortex BA6 R<br>HOCSA:<br>36% Precentral Gyrus, 22% Cingulate Gyrus, posterior division, 12% Postcentral Gyrus, 8% Precuneous Cortex |
| 6b | -6.932 | -35.468 | 50.75 | 4.557 | | JHA:<br>44% GM Superior parietal lobule 5M L, 21% GM Primary motor cortex BA4a L, 10% GM Superior parietal lobule 5Ci L, 6% WM Corticospinal tract L<br>HOCSA:<br>29% Precentral Gyrus, 21% Precuneous Cortex, 15% Postcentral Gyrus, 12% Cingulate Gyrus, posterior division |
| 6c | -14.396 | -32.98 | 39.75 | 4.473 | | JHA:<br>46% GM Superior parietal lobule 5Ci L, 2% GM Superior parietal lobule 5M L<br>HOCSA:<br>29% Cingulate Gyrus, posterior division, 29% Precentral Gyrus, 11% Precuneous Cortex, 2% Postcentral Gyrus |
| 7 | 27.899 | -8.1 | -18.0 | 5.155 | 1787 | JHA:<br>79% GM Amygdala_laterobasal group R, 74% GM Hippocampus cornu ammonis R, 15% GM Hippocampus dentate gyrus R, 5% GM Amygdala_superficial group R, 3% GM Hippocampus subiculum R, 1% GM Amygdala_centromedial group R |
| 8 | 52.779 | 1.851 | -34.5 | 4.946 | 987 | HOCSA:<br>40% Middle Temporal Gyrus, anterior division, 14% Inferior Temporal Gyrus, anterior division, 6% Temporal Pole, 2% Middle Temporal Gyrus, posterior division |
| 9 | -59.18 | -28.0 | 15.0 | 4.882 | 1225 | JHA:<br>36% GM Secondary somatosensory cortex / Parietal operculum OP1 L, 24% GM Inferior parietal lobule PFop L, 2% GM Inferior parietal lobule PFcm L, 2% GM Inferior parietal lobule PF L<br>HOCSA:<br>46% Parietal Operculum Cortex, 30% Planum Temporale, 4% Central Opercular Cortex, 4% Supramarginal Gyrus, anterior division, 1% Superior Temporal Gyrus, posterior division |
| 10 | -24.348 | 49.123 | 37.0 | 4.663 | 561 | HOCSA:<br>77% Frontal Pole |
| 11 | -6.932 | 36.683 | -4.25 | 4.337 | 1004 | HOCSA:<br>56% Cingulate Gyrus, anterior division, 17% Paracingulate Gyrus, 1% Subcallosal Cortex, 1% Frontal Medial Cortex |
| 11a | 3.019 | 29.219 | -7.0 | 4.221 | | HOCSA:<br>56% Subcallosal Cortex, 18% Cingulate Gyrus, anterior division, 6% Paracingulate Gyrus, 1% Frontal Medial Cortex |
| 12 | 22.923 | -42.932 | 56.25 | 4.302 | 442 | JHA:<br>77% GM Primary somatosensory cortex BA2 R, 37% GM Superior parietal lobule 7PC R, 26% GM Superior parietal lobule |

| | | | | | | |
|---|---|---|---|---|---|---|
| | | | | | | 5L R, 15% GM Primary somatosensory cortex BA1 R, 10% GM Primary motor cortex BA4p R, 9% GM Primary somatosensory cortex BA3b R, 6% GM Superior parietal lobule 5M R, 1% GM Anterior intra-parietal sulcus hIP3 R<br>HOCSA:<br>22% Postcentral Gyrus, 20% Superior Parietal Lobule |
| 12a | 25.411 | -40.444 | 64.5 | 3.749 | | JHA:<br>48% GM Primary somatosensory cortex BA2 R, 30% GM Primary somatosensory cortex BA3b R, 23% GM Primary somatosensory cortex BA1 R, 10% GM Superior parietal lobule 7PC R, 8% GM Primary motor cortex BA4a R, 6% GM Superior parietal lobule 7A R, 6% GM Primary motor cortex BA4p R, 4% GM Superior parietal lobule 5M R, 4% GM Superior parietal lobule 5L R, 2% GM Premotor cortex BA6 R, 1% WM Corticospinal tract R<br>HOCSA:<br>35% Superior Parietal Lobule, 28% Postcentral Gyrus |
| 13 | 40.339 | -15.564 | 20.5 | 4.267 | 731 | JHA:<br>61% GM Secondary somatosensory cortex / Parietal operculum OP3 R, 53% GM Secondary somatosensory cortex / Parietal operculum OP2 R, 13% GM Insula Ig2 R, 13% GM Secondary somatosensory cortex / Parietal operculum OP4 R, 10% GM Secondary somatosensory cortex / Parietal operculum OP1 R<br><br>HOCSA:<br>60% Central Opercular Cortex, 15% Parietal Operculum Cortex, 3% Insular Cortex\n' |
| 13a | 50.291 | -15.564 | 28.75 | 3.889 | | JHA:<br>36% GM Inferior parietal lobule PFop R, 15% GM Inferior parietal lobule PFt R, 10% GM Secondary somatosensory cortex / Parietal operculum OP4 R, 7% GM Secondary somatosensory cortex / Parietal operculum OP3 R, 6% GM Secondary somatosensory cortex / Parietal operculum OP1 R, 6% GM Primary somatosensory cortex BA3b R, 1% GM Primary somatosensory cortex BA3a R<br>HOCSA:<br>10% Postcentral Gyrus, 3% Supramarginal Gyrus, anterior division |
| 14 | 40.339 | -35.468 | 15.0 | 4.225 | 204 | JHA:<br>4% WM Optic radiation R<br>HOCSA:<br>25% Planum Temporale, 6% Supramarginal Gyrus, posterior division, 3% Parietal Operculum Cortex |
| 15 | 42.827 | -42.932 | -18.0 | 4.159 | 238 | HOCSA:<br>34% Temporal Occipital Fusiform Cortex, 17% Inferior Temporal Gyrus, temporooccipital part, 10% Temporal Fusiform Cortex, posterior division, 1% Inferior Temporal Gyrus, posterior division |
| 16 | 42.827 | -10.588 | 53.5 | 4.094 | 255 | JHA:<br>48% GM Premotor cortex BA6 R, 22% GM Primary motor cortex BA4a R, 16% WM Corticospinal tract R, 10% GM Primary somatosensory cortex BA1 R, 6% GM Primary somatosensory cortex BA3b R<br>HOCSA:<br>48% Precentral Gyrus, 2% Postcentral Gyrus |
| 17 | 65.219 | -37.956 | 1.25 | 3.938 | 204 | HOCSA:<br>36% Middle Temporal Gyrus, temporooccipital part, 30% Middle Temporal Gyrus, posterior division, 12% Supramarginal Gyrus, posterior division, 7% Superior Temporal Gyrus, posterior division |
| 18 | 35.363 | 31.707 | 48.0 | 3.846 | 221 | HOCSA:<br>51% Middle Frontal Gyrus, 7% Frontal Pole, 2% Superior Frontal Gyrus |
| 19 | -24.348 | 56.587 | 23.25 | 3.814 | 221 | HOCSA: 81% Frontal Pole |

**Supplementary Table 3: Significant clusters of the action similarity-based BOLD adaptation analysis of the action combinations.** Clusters surviving the whole-brain correction using FDR method with a voxel-level threshold of p<0.01. Table displays MNI coordinates (X, Y, Z), statistical T values and atlas labels of peak voxels of the clusters. Atlas labels are based on the Juelich Histological Atlas (JHA) and Harvard-Oxford Cortical Structural Atlas (HOCSA). The labels were generated using FSL. Atlases are listed only if the labels were found for a given atlas. The letters after the cluster ID indicates its subcluster.

| Cluster ID | X | Y | Z | T Value | Cluster Size (voxels) | Atlas label |
|---|---|---|---|---|---|---|
| 1 | -56.692 | -52.884 | 6.75 | -7.433 | 16733 | HOCSA:<br>54% Middle Temporal Gyrus, temporooccipital part, 9% Angular Gyrus, 7% Supramarginal Gyrus, posterior division, 3% Middle Temporal Gyrus, posterior division, 1% Lateral Occipital Cortex, inferior division |
| 1a | -66.644 | -30.492 | 4.0 | -6.462 | | JHA:<br>2% GM Secondary somatosensory cortex / Parietal operculum OP1 L, 2% GM Inferior parietal lobule PF L<br><br>HOCSA:<br>69% Superior Temporal Gyrus, posterior division, 8% Middle Temporal Gyrus, posterior division, 1% Planum Temporale |
| 1b | -61.668 | -60.348 | 6.75 | -6.381 | | JHA:<br>3% GM Inferior parietal lobule Pga L<br><br>HOCSA:<br>47% Middle Temporal Gyrus, temporooccipital part, 20% Lateral Occipital Cortex, inferior division, 6% Angular Gyrus, 2% Lateral Occipital Cortex, superior division |
| 1c | -59.18 | -42.932 | 23.25 | -6.26 | | JHA:<br>70% GM Inferior parietal lobule PF L, 35% GM Inferior parietal lobule PFm L, 14% GM Inferior parietal lobule PFcm L<br><br>HOCSA:<br>36% Supramarginal Gyrus, posterior division, 22% Parietal Operculum Cortex, 11% Planum Temporale, 4% Supramarginal Gyrus, anterior division, 4% Superior Temporal Gyrus, posterior division, 2% Angular Gyrus |
| 2 | -49.228 | -75.276 | -9.75 | -6.38 | 5992 | JHA:<br>4% GM Visual cortex V5 L<br><br>HOCSA:<br>77% Lateral Occipital Cortex, inferior division, 1% Occipital Fusiform Gyrus |
| 2a | -41.764 | -80.252 | -9.75 | -5.583 | | JHA:<br>7% GM Visual cortex V4 L<br><br>HOCSA:<br>67% Lateral Occipital Cortex, inferior division, 4% Occipital Fusiform Gyrus |
| 2b | -44.252 | -85.228 | 1.25 | -4.945 | | JHA:<br>14% GM Visual cortex V4 L, HOCSA: 63% Lateral Occipital Cortex, inferior division, 6% Occipital Pole, 2% Lateral Occipital Cortex, superior division |
| 2c | -49.228 | -60.348 | -18.0 | -4.128 | | HOCSA:<br>47% Inferior Temporal Gyrus, temporooccipital part, 18% Lateral Occipital Cortex, inferior division, 14% Temporal Occipital Fusiform Cortex, 3% Occipital Fusiform Gyrus, 2% Middle Temporal Gyrus, temporooccipital part |
| 3 | 47.803 | -80.252 | 6.75 | -6.125 | 16971 | JHA:<br>14% GM Visual cortex V5 R, 2% GM Inferior parietal lobule PGp R, 1% GM Visual cortex V4 R<br><br>HOCSA:<br>70% Lateral Occipital Cortex, inferior division, 5% Lateral Occipital Cortex, superior division |

| | | | | | | |
|---|---|---|---|---|---|---|
| 3a | 57.755 | -30.492 | 37.0 | -5.98 | | JHA:<br>52% GM Inferior parietal lobule PF R, 22% GM Inferior parietal lobule PFop R, 18% GM Inferior parietal lobule PFcm R, 10% GM Anterior intra-parietal sulcus hIP2 R, 8% GM Secondary somatosensory cortex / Parietal operculum OP1 R, 8% GM Inferior parietal lobule PFt R, 5% GM Inferior parietal lobule PFm R<br><br>HOCSA:<br>51% Supramarginal Gyrus, anterior division, 9% Parietal Operculum Cortex, 7% Supramarginal Gyrus, posterior division, 4% Postcentral Gyrus, 2% Planum Temporale |
| 3b | 62.731 | -55.372 | 9.5 | -5.676 | | JHA:<br>7% GM Inferior parietal lobule Pga R<br><br>HOCSA:<br>53% Middle Temporal Gyrus, temporooccipital part, 15% Angular Gyrus, 7% Lateral Occipital Cortex, inferior division, 2% Lateral Occipital Cortex, superior division |
| 3c | 55.267 | -42.932 | 9.5 | -5.547 | | JHA:<br>9% GM Inferior parietal lobule PFm R, 3% GM Inferior parietal lobule Pga R<br><br>HOCSA:<br>36% Supramarginal Gyrus, posterior division, 22% Middle Temporal Gyrus, temporooccipital part, 10% Angular Gyrus, 4% Superior Temporal Gyrus, posterior division, 1% Middle Temporal Gyrus, posterior division |
| 4 | -59.18 | 14.291 | 12.25 | -5.535 | 3251 | JHA:<br>48% GM Broca's area BA44 L, 16% GM Broca's area BA45 L, 1% GM Primary somatosensory cortex BA3b L<br><br>HOCSA:<br>27% Inferior Frontal Gyrus, pars opercularis, 7% Precentral Gyrus |
| 4a | -61.668 | 19.267 | 26.0 | -4.76 | | JHA:<br>5% GM Broca's area BA45 L, 4% GM Broca's area BA44 L |
| 4b | -51.716 | 16.779 | 20.5 | -4.672 | | JHA:<br>44% GM Broca's area BA44 L, 13% GM Broca's area BA45 L<br><br>HOCSA:<br>58% Inferior Frontal Gyrus, pars opercularis, 5% Precentral Gyrus, 1% Inferior Frontal Gyrus, pars triangularis |
| 4c | -49.228 | 9.315 | 9.5 | -3.795 | | JHA:<br>27% GM Broca's area BA44 L, 2% GM Broca's area BA45 L<br><br>HOCSA:<br>45% Inferior Frontal Gyrus, pars opercularis, 11% Precentral Gyrus, 1% Frontal Operculum Cortex |
| 5 | -54.204 | -0.636 | 42.5 | -5.496 | 3557 | JHA:<br>68% GM Premotor cortex BA6 L, 4% GM Broca's area BA44 L, 1% WM Corticospinal tract L, 1% GM Primary somatosensory cortex BA1 L, 1% GM Primary motor cortex BA4a L<br><br>HOCSA:<br>75% Precentral Gyrus, 8% Middle Frontal Gyrus |
| 5a | -41.764 | 1.851 | 50.75 | -5.085 | | JHA:<br>24% GM Premotor cortex BA6 L<br><br>HOCSA:<br>36% Middle Frontal Gyrus, 26% Precentral Gyrus |
| 5b | -41.764 | -10.588 | 42.5 | -4.011 | | JHA:<br>38% WM Corticospinal tract L, 38% GM Primary motor cortex BA4a L, 33% GM Primary motor cortex BA4p L, 11% GM Primary somatosensory cortex BA3b L, 9% GM Premotor cortex BA6 L, 1% GM Primary somatosensory cortex BA1 L |

| | | | | | | |
|---|---|---|---|---|---|---|
| | | | | | | HOCSA:<br>30% Precentral Gyrus, 1% Postcentral Gyrus |
| 6 | 42.827 | -52.884 | -15.25 | -5.323 | 5055 | HOCSA:<br>53% Temporal Occipital Fusiform Cortex, 10% Inferior Temporal Gyrus, temporooccipital part |
| 6a | 32.875 | -70.3 | -15.25 | -5.122 | | JHA:<br>48% GM Visual cortex V4 R, 3% GM Visual cortex V3V R<br><br>HOCSA:<br>75% Occipital Fusiform Gyrus, 2% Temporal Occipital Fusiform Cortex, 1% Lingual Gyrus, 1% Lateral Occipital Cortex, inferior division |
| 6b | 42.827 | -65.324 | -15.25 | -4.949 | | JHA:<br>8% GM Visual cortex V4 R<br><br>HOCSA:<br>43% Occipital Fusiform Gyrus, 21% Lateral Occipital Cortex, inferior division, 4% Temporal Occipital Fusiform Cortex, 3% Inferior Temporal Gyrus, temporooccipital part |
| 6c | 27.899 | -50.396 | -9.75 | -4.693 | | JHA:<br>1% WM Optic radiation R<br><br>HOCSA: 54% Temporal Occipital Fusiform Cortex, 20% Lingual Gyrus |
| 7 | -34.3 | 56.587 | 20.5 | -5.096 | 476 | HOCSA:<br>72% Frontal Pole |
| 8 | -36.788 | 41.659 | 37.0 | -4.797 | 1174 | HOCSA:<br>51% Frontal Pole, 16% Middle Frontal Gyrus |
| 9 | -6.932 | 4.339 | 67.25 | -4.673 | 1957 | JHA:<br>73% GM Premotor cortex BA6 L<br><br>HOCSA:<br>23% Juxtapositional Lobule Cortex (formerly Supplementary Motor Cortex), 20% Superior Frontal Gyrus |
| 9a | -19.372 | 11.803 | 67.25 | -4.132 | | JHA:<br>16% GM Premotor cortex BA6 L<br><br>HOCSA:<br>42% Superior Frontal Gyrus, 1% Middle Frontal Gyrus |
| 10 | 17.947 | -97.668 | -1.5 | -4.662 | 629 | JHA:<br>88% GM Visual cortex V1 BA17 R, 51% WM Optic radiation R, 25% GM Visual cortex V2 BA18 R<br><br>HOCSA:<br>62% Occipital Pole, 1% Lateral Occipital Cortex, inferior division |
| 10a | 15.459 | -102.644 | 6.75 | -4.081 | | JHA:<br>78% GM Visual cortex V1 BA17 R, 38% GM Visual cortex V2 BA18 R, 26% WM Optic radiation R<br><br>HOCSA:<br>71% Occipital Pole |
| 11 | 60.243 | 9.315 | 12.25 | -4.513 | 391 | JHA:<br>53% GM Broca's area BA44 R, 6% GM Secondary somatosensory cortex / Parietal operculum OP4 R, 2% GM Broca's area BA45 R<br><br>HOCSA:<br>38% Precentral Gyrus, 28% Inferior Frontal Gyrus, pars opercularis |
| 12 | -14.396 | -75.276 | -48.25 | -4.459 | 680 | Could not be labeled |
| 13 | 25.411 | 1.851 | 31.5 | -4.356 | 306 | JHA:<br>3% WM Callosal body |
| 14 | -44.252 | -42.932 | -18.0 | -4.328 | 340 | HOCSA:<br>26% Temporal Fusiform Cortex, posterior division, 21% Inferior Temporal Gyrus, posterior division, 10% Inferior Temporal Gyrus, temporooccipital part, 6% Temporal Occipital Fusiform Cortex |

| | | | | | | |
|---|---|---|---|---|---|---|
| 15 | 37.851 | -87.716 | -9.75 | -4.191 | 272 | JHA:<br>44% GM Visual cortex V3V R, 36% GM Visual cortex V4 R, 22% GM Visual cortex V2 BA18 R<br><br>HOCSA:<br>51% Lateral Occipital Cortex, inferior division, 21% Occipital Pole, 2% Occipital Fusiform Gyrus |
| 16 | 47.803 | -18.052 | -7.0 | -4.105 | 306 | JHA:<br>11% GM Insula Id1 R<br><br>HOCSA:<br>47% Superior Temporal Gyrus, posterior division, 19% Middle Temporal Gyrus, posterior division, 1% Middle Temporal Gyrus, anterior division, 1% Superior Temporal Gyrus, anterior division |
| 17 | -61.668 | 1.851 | 26.0 | -3.822 | 238 | JHA:<br>48% GM Premotor cortex BA6 L, 19% GM Broca's area BA44 L, 10% GM Primary somatosensory cortex BA3b L, 6% GM Primary somatosensory cortex BA1 L<br><br>HOCSA:<br>52% Precentral Gyrus, 3% Postcentral Gyrus |

**Supplementary Table 4: Whole-brain clusters of the generalized Psychophysiological Interaction (gPPI) effect with left HPC used as a seed region.** No clusters survived FDR correction in the whole-brain analysis using a voxel-level threshold of $p<0.01$. The table lists clusters uncorrected for multiple comparisons, with statistical significance threshold defined at a single voxel level of $z_\alpha=1.96$. Table displays MNI coordinates (X, Y, Z), statistical T values and atlas labels of peak voxels of the clusters. Atlas labels are based on the Juelich Histological Atlas (JHA) and Harvard-Oxford Cortical Structural Atlas (HOCSA). The labels were generated using FSL. Atlases are listed only if the labels were found for a given atlas. The letters after the cluster ID indicates its subcluster.

| Cluster ID | X | Y | Z | T Value | Cluster Size (voxels) | Atlas label |
|---|---|---|---|---|---|---|
| 1 | -29.324 | 49.123 | -12.5 | 4.41 | 783 | HOCSA:<br>69% Frontal Pole |
| 2 | -41.764 | 46.635 | 28.75 | 4.095 | 1651 | HOCSA:<br>34% Frontal Pole |
| 2a | -51.716 | 41.659 | 23.25 | 2.972 | | HOCSA:<br>1% Frontal Pole |
| 3 | 10.483 | -32.98 | 42.5 | 4.045 | 13890 | JHA:<br>39% GM Superior parietal lobule 5Ci R, 18% GM Premotor cortex BA6 R, 1% GM Superior parietal lobule 5M R, 1% GM Primary motor cortex BA4a R<br><br>HOCSA:<br>60% Cingulate Gyrus, posterior division, 9% Precuneous Cortex, 3% Postcentral Gyrus, 3% Precentral Gyrus |
| 3a | 7.995 | -32.98 | 28.75 | 3.938 | | JHA:<br>50% WM Callosal body<br><br>HOCSA:<br>35% Cingulate Gyrus, posterior division |
| 3b | 5.507 | -37.956 | 53.5 | 3.876 | | JHA:<br>79% GM Superior parietal lobule 5M R, 32% GM Primary motor cortex BA4a R, 20% GM Superior parietal lobule 5Ci R, 11% WM Corticospinal tract R<br><br>HOCSA:<br>29% Precuneous Cortex, 29% Postcentral Gyrus, 10% Precentral Gyrus, 4% Cingulate Gyrus, posterior division |

| | | | | | | |
|---|---|---|---|---|---|---|
| 3c | -4.444 | -52.884 | 67.25 | 3.716 | | JHA:<br>40% GM Superior parietal lobule 5L L, 32% GM Superior parietal lobule 7A L, 22% GM Primary somatosensory cortex BA3b L, 14% GM Superior parietal lobule 7PC L, 11% GM Superior parietal lobule 5M L, 4% GM Primary motor cortex BA4a L, 3% GM Primary motor cortex BA4p L<br><br>HOCSA:<br>41% Precuneous Cortex, 7% Postcentral Gyrus, 6% Superior Parietal Lobule, 2% Lateral Occipital Cortex, superior division |
| 4 | 37.851 | -23.028 | -20.75 | 3.972 | 1072 | JHA:<br>34% GM Hippocampus cornu ammonis R, 14% WM Optic radiation R, 11% GM Hippocampus dentate gyrus R, 2% GM Hippocampus subiculum R<br><br>HOCSA:<br>38% Temporal Fusiform Cortex, posterior division, 5% Parahippocampal Gyrus, posterior division, 4% Parahippocampal Gyrus, anterior division, 2% Inferior Temporal Gyrus, posterior division, 1% Temporal Occipital Fusiform Cortex |
| 5 | -14.396 | -23.028 | 15.0 | 3.843 | 919 | JHA:<br>1% WM Corticospinal tract L |
| 5a | -16.884 | -13.076 | 9.5 | 2.522 | | JHA:<br>37% WM Corticospinal tract L |
| 6 | -11.908 | -40.444 | 83.75 | 3.795 | 1566 | JHA:<br>4% GM Superior parietal lobule 5M L HOCSA: 2% Postcentral Gyrus |
| 6a | -14.396 | -52.884 | 78.25 | 3.556 | | JHA:<br>9% GM Superior parietal lobule 7PC L, 9% GM Superior parietal lobule 7A L, 6% GM Superior parietal lobule 5L L, 5% GM Primary motor cortex BA4a L, 2% GM Superior parietal lobule 5M L<br><br>HOCSA:<br>9% Superior Parietal Lobule, 7% Postcentral Gyrus, 1% Lateral Occipital Cortex, superior division |
| 6b | -11.908 | -45.42 | 81.0 | 2.688 | | JHA:<br>41% GM Superior parietal lobule 5L L, 23% GM Superior parietal lobule 5M L, 19% GM Primary somatosensory cortex BA1 L, 11% GM Primary somatosensory cortex BA3b L, 11% GM Primary motor cortex BA4a L, 9% GM Primary motor cortex BA4p L, 4% GM Premotor cortex BA6 L<br><br>HOCSA:<br>17% Postcentral Gyrus, 4% Superior Parietal Lobule |
| 6c | -26.836 | -40.444 | 72.75 | 2.436 | | JHA:<br>53% GM Primary somatosensory cortex BA1 L, 24% GM Primary somatosensory cortex BA2 L, 23% GM Superior parietal lobule 5L L, 16% GM Superior parietal lobule 7PC L, 16% GM Superior parietal lobule 7A L, 13% GM Primary motor cortex BA4a L, 8% GM Primary motor cortex BA4p L, 6% GM Primary somatosensory cortex BA3b L, 1% WM Corticospinal tract L<br><br>HOCSA:<br>33% Postcentral Gyrus, 11% Superior Parietal Lobule, 3% Supramarginal Gyrus, anterior division |
| 7 | 0.531 | 19.267 | 34.25 | 3.756 | 7813 | HOCSA:<br>58% Cingulate Gyrus, anterior division, 22% Paracingulate Gyrus |
| 7a | 10.483 | -3.124 | 53.5 | 3.708 | | JHA:<br>45% GM Premotor cortex BA6 R<br><br>HOCSA:<br>34% Juxtapositional Lobule Cortex (formerly Supplementary Motor Cortex), 3% Precentral Gyrus, 1% Cingulate Gyrus, anterior division |
| 7b | -9.42 | -3.124 | 39.75 | 3.414 | | JHA:<br>1% WM Cingulum L, 1% GM Premotor cortex BA6 L<br><br>HOCSA: |

| | | | | | | |
|---|---|---|---|---|---|---|
| | | | | | | 22% Juxtapositional Lobule Cortex (formerly Supplementary Motor Cortex), 19% Cingulate Gyrus, anterior division |
| 7c | 3.019 | 1.851 | 48.0 | 3.175 | | JHA:<br>44% GM Premotor cortex BA6 R<br><br>HOCSA:<br>58% Juxtapositional Lobule Cortex (formerly Supplementary Motor Cortex), 24% Cingulate Gyrus, anterior division, 3% Paracingulate Gyrus |
| 8 | 35.363 | 14.291 | -1.5 | 3.75 | 408 | JHA:<br>17% WM Inferior occipito-frontal fascicle R<br><br>HOCSA:<br>61% Insular Cortex |
| 9 | 40.339 | 46.635 | 28.75 | 3.568 | 1889 | HOCSA:<br>85% Frontal Pole, 1% Middle Frontal Gyrus |
| 9a | 27.899 | 59.075 | 28.75 | 2.523 | | HOCSA:<br>35% Frontal Pole |
| 10 | 15.459 | -8.1 | 4.0 | 3.55 | 2161 | JHA:<br>25% WM Corticospinal tract R |
| 10a | 7.995 | -10.588 | 15.0 | 3.32 | | JHA:<br>11% WM Fornix |
| 10b | 12.971 | -20.54 | 12.25 | 2.31 | | JHA:<br>1% WM Fornix |
| 11 | 65.219 | -10.588 | 42.5 | 3.536 | 187 | HOCSA:<br>2% Postcentral Gyrus |
| 12 | 47.803 | -77.764 | 37.0 | 3.504 | 1395 | JHA:<br>3% GM Inferior parietal lobule PGp R<br><br>HOCSA:<br>12% Lateral Occipital Cortex, superior division |
| 12a | 42.827 | -65.324 | 28.75 | 2.887 | | JHA:<br>50% GM Inferior parietal lobule PGp R<br><br>HOCSA:<br>46% Lateral Occipital Cortex, superior division, 1% Angular Gyrus |
| 12b | 47.803 | -75.276 | 26.0 | 2.432 | | JHA:<br>78% GM Inferior parietal lobule PGp R<br><br>HOCSA:<br>73% Lateral Occipital Cortex, superior division, 1% Lateral Occipital Cortex, inferior division |
| 13 | -36.788 | 4.339 | 17.75 | 3.493 | 766 | JHA:<br>6% GM Broca's area BA44 L<br><br>HOCSA:<br>7% Central Opercular Cortex |
| 14 | 52.779 | -35.468 | 31.5 | 3.467 | 4528 | JHA:<br>23% GM Inferior parietal lobule PFm R, 21% GM Inferior parietal lobule PF R, 20% GM Inferior parietal lobule PFcm R, 12% GM Anterior intra-parietal sulcus hIP2 R, 8% GM Anterior intra-parietal sulcus hIP1 R<br><br>HOCSA:<br>16% Parietal Operculum Cortex, 15% Supramarginal Gyrus, posterior division, 5% Supramarginal Gyrus, anterior division, 4% Planum Temporale, 1% Angular Gyrus |
| 14a | 62.731 | -30.492 | 28.75 | 3.251 | | JHA:<br>73% GM Inferior parietal lobule PF R, 45% GM Inferior parietal lobule PFcm R, 3% GM Inferior parietal lobule PFop R, 2% GM Secondary somatosensory cortex / Parietal operculum OP1 R<br><br>HOCSA:<br>40% Supramarginal Gyrus, anterior division, 20% Parietal Operculum Cortex, 13% Planum Temporale, 4% Supramarginal Gyrus, posterior division, 1% Superior Temporal Gyrus, posterior division |
| 14b | 62.731 | -40.444 | 31.5 | 2.987 | | JHA: |

| | | | | | | |
|---|---|---|---|---|---|---|
| | | | | | | 83% GM Inferior parietal lobule PF R, 50% GM Inferior parietal lobule PFm R, 1% GM Inferior parietal lobule Pga R<br><br>HOCSA:<br>58% Supramarginal Gyrus, posterior division, 10% Angular Gyrus, 4% Planum Temporale, 2% Supramarginal Gyrus, anterior division |
| 14c | 65.219 | -50.396 | 34.25 | 2.859 | | HOCSA:<br>13% Angular Gyrus, 1% Supramarginal Gyrus, posterior divisio |
| 15 | 35.363 | 26.73 | -20.75 | 3.372 | 442 | HOCSA:<br>75% Frontal Orbital Cortex |
| 16 | -71.62 | -28 | 26.0 | 3.316 | 1225 | No label found |
| 16a | -61.668 | -35.468 | 26.0 | 2.734 | | JHA:<br>80% GM Inferior parietal lobule PF L, 6% GM Inferior parietal lobule PFcm L, 4% GM Anterior intra-parietal sulcus hIP2 L, 3% GM Inferior parietal lobule PFm L<br><br>HOCSA:<br>41% Supramarginal Gyrus, anterior division, 21% Parietal Operculum Cortex, 13% Planum Temporale, 5% Supramarginal Gyrus, posterior division |
| 17 | -29.324 | 6.827 | -29.0 | 3.241 | 323 | HOCSA:<br>73% Temporal Pole, 3% Parahippocampal Gyrus, anterior division |
| 18 | -41.764 | -32.98 | -45.5 | 3.231 | 544 | No label found |
| 18a | -41.764 | -45.42 | -51.0 | 2.325 | | No label found |
| 19 | -11.908 | -0.636 | -1.5 | 3.185 | 510 | No label found |
| 20 | -39.276 | -18.052 | 28.75 | 3.137 | 561 | JHA:<br>8% GM Primary somatosensory cortex BA3a L, 2% GM Secondary somatosensory cortex / Parietal operculum OP3 L, 1% GM Insula Ig2 L, 1% GM Secondary somatosensory cortex / Parietal operculum OP1 L<br><br>HOCSA:<br>2% Postcentral Gyrus |
| 20a | -29.324 | -20.54 | 31.5 | 2.623 | | JHA:<br>50% WM Corticospinal tract L, 23% WM Superior longitudinal fascicle L |
| 20b | -31.812 | -18.052 | 23.25 | 2.423 | | JHA:<br>25% WM Superior longitudinal fascicle L, 14% GM Secondary somatosensory cortex / Parietal operculum OP3 L, 11% GM Secondary somatosensory cortex / Parietal operculum OP2 L, 9% WM Corticospinal tract L, 5% GM Insula Ig2 L<br><br>HOCSA:<br>1% Central Opercular Cortex |
| 21 | 27.899 | -23.028 | -1.5 | 3.044 | 272 | JHA:<br>67% WM Optic radiation R, 15% WM Acoustic radiation R, 2% GM Lateral geniculate body R, 1% WM Corticospinal tract R |
| 22 | 37.851 | -8.1 | -4.25 | 3.032 | 1021 | JHA:<br>12% WM Inferior occipito-frontal fascicle R, 8% GM Insula Id1 R, 2% WM Acoustic radiation R, 1% GM Insula Ig2 R<br><br>HOCSA:<br>47% Insular Cortex |
| 22a | 37.851 | -3.124 | -15.25 | 2.153 | | JHA:<br>24% WM Optic radiation R, 23% WM Inferior occipito-frontal fascicle R, 14% WM Uncinate fascicle R, 4% GM Insula Id1 R, 4% GM Amygdala_superficial group R, 4% GM Amygdala_centromedial group R, 2% GM Amygdala_laterobasal group R<br><br>HOCSA:<br>13% Insular Cortex, 4% Planum Polare |
| 23 | -66.644 | -37.956 | 42.5 | 3.017 | 714 | HOCSA:<br>8% Supramarginal Gyrus, anterior division, 4% Supramarginal Gyrus, posterior division, 1% Planum Temporale, 1% Parietal Operculum Cortex |
| 23a | -64.156 | -30.492 | 48.0 | 2.633 | | HOCSA:<br>12% Supramarginal Gyrus, anterior division, 1% Postcentral Gyrus |

| | | | | | | |
|---|---|---|---|---|---|---|
| 24 | 47.803 | 19.267 | -18.0 | 2.974 | 425 | HOCSA:<br>67% Temporal Pole, 6% Frontal Orbital Cortex |
| 25 | 10.483 | -15.564 | 81.0 | 2.945 | 425 | JHA:<br>51% GM Premotor cortex BA6 R, 6% GM Primary motor cortex BA4a R<br><br>HOCSA:<br>24% Precentral Gyrus, 4% Superior Frontal Gyrus |
| 25a | 17.947 | -13.076 | 78.25 | 2.263 | | JHA:<br>98% GM Premotor cortex BA6 R<br><br>HOCSA:<br>24% Precentral Gyrus, 16% Superior Frontal Gyrus |
| 26 | 55.267 | 16.779 | -4.25 | 2.939 | 1174 | JHA:<br>2% GM Broca's area BA45 R<br><br>HOCSA:<br>16% Temporal Pole, 11% Inferior Frontal Gyrus, pars opercularis, 2% Frontal Operculum Cortex, 2% Frontal Orbital Cortex, 2% Precentral Gyrus, 2% Inferior Frontal Gyrus, pars triangularis, 1% Central Opercular Cortex |
| 27 | 70.195 | -13.076 | 26.0 | 2.926 | 493 | JHA:<br>1% GM Secondary somatosensory cortex / Parietal operculum OP4 R, 1% GM Secondary somatosensory cortex / Parietal operculum OP1 R, 1% GM Inferior parietal lobule PFt R, 1% GM Inferior parietal lobule PFop R, 1% GM Inferior parietal lobule PF R<br><br>HOCSA:<br>4% Postcentral Gyrus, 1% Supramarginal Gyrus, anterior division |
| 27a | 67.707 | -10.588 | 37.0 | 2.754 | | HOCSA:<br>1% Postcentral Gyrus |
| 28 | -11.908 | -87.716 | 48.0 | 2.898 | 714 | JHA:<br>11% GM Superior parietal lobule 7P L, 8% GM Superior parietal lobule 7A L<br><br>HOCSA:<br>10% Lateral Occipital Cortex, superior division, 9% Occipital Pole |
| 29 | -14.396 | -0.636 | 75.5 | 2.888 | 476 | JHA:<br>41% GM Premotor cortex BA6 L<br><br>HOCSA:<br>22% Superior Frontal Gyrus |
| 30 | 32.875 | -0.636 | 28.75 | 2.859 | 578 | JHA:<br>7% WM Corticospinal tract R |
| 30a | 25.411 | -10.588 | 31.5 | 2.503 | | JHA:<br>42% WM Corticospinal tract R, 1% WM Superior occipito-frontal fascicle R |
| 31 | 12.971 | 9.315 | 4.0 | 2.849 | 459 | No label found |
| 32 | 0.531 | -47.908 | -73.0 | 2.82 | 425 | No label found |
| 33 | -24.348 | 29.219 | 9.5 | 2.763 | 306 | JHA:<br>12% WM Callosal body, 8% WM Superior occipito-frontal fascicle L<br><br>HOCSA:<br>1% Frontal Orbital Cortex |
| 34 | 45.315 | -23.028 | 64.5 | 2.735 | 255 | JHA:<br>70% GM Primary somatosensory cortex BA1 R, 6% GM Premotor cortex BA6 R, 6% GM Primary somatosensory cortex BA3b R, 4% GM Primary somatosensory cortex BA2 R, 4% GM Primary motor cortex BA4a R<br><br>HOCSA:<br>63% Postcentral Gyrus, 2% Precentral Gyrus |
| 35 | -16.884 | 31.707 | -15.25 | 2.695 | 442 | JHA:<br>3% WM Callosal body<br><br>HOCSA:<br>5% Frontal Orbital Cortex, 3% Frontal Pole, 2% Frontal Medial Cortex |

| | | | | | | |
|---|---|---|---|---|---|---|
| 35a | -14.396 | 21.755 | -7.0 | 2.515 | | JHA:<br>27% WM Callosal body |
| 36 | -4.444 | -35.468 | -56.5 | 2.674 | 289 | No label found |
| 36a | -11.908 | -37.956 | -59.25 | 2.171 | | No label found |
| 37 | -24.348 | 6.827 | 26.0 | 2.654 | 340 | JHA:<br>15% WM Superior occipito-frontal fascicle L |
| 38 | -31.812 | 31.707 | -1.5 | 2.62 | 374 | HOCSA:<br>31% Frontal Orbital Cortex, 3% Inferior Frontal Gyrus, pars triangularis, 1% Frontal Pole |
| 39 | -39.276 | -13.076 | -12.5 | 2.594 | 255 | JHA:<br>20% WM Optic radiation L, 18% WM Inferior occipito-frontal fascicle L, 8% GM Insula Id1 L<br><br>HOCSA:<br>18% Planum Polare, 1% Insular Cortex |
| 40 | -26.836 | -97.668 | -9.75 | 2.557 | 238 | JHA:<br>47% GM Visual cortex V2 BA18 L, 35% GM Visual cortex V3V L, 22% GM Visual cortex V1 BA17 L, 7% WM Optic radiation L, 3% GM Visual cortex V4 L<br><br>HOCSA:<br>62% Occipital Pole, 4% Lateral Occipital Cortex, inferior division |
| 41 | 30.387 | 51.611 | -12.5 | 2.546 | 510 | HOCSA:<br>63% Frontal Pole |
| 42 | 32.875 | 39.171 | 48.0 | 2.529 | 646 | HOCSA:<br>15% Frontal Pole, 4% Middle Frontal Gyrus |
| 42a | 30.387 | 31.707 | 42.5 | 2.387 | | HOCSA:<br>48% Middle Frontal Gyrus, 10% Frontal Pole, 8% Superior Frontal Gyrus |
| 43 | 27.899 | -95.18 | -7.0 | 2.518 | 221 | JHA:<br>76% GM Visual cortex V2 BA18 R, 43% GM Visual cortex V1 BA17 R, 23% WM Optic radiation R, 20% GM Visual cortex V3V R<br><br>HOCSA:<br>58% Occipital Pole, 6% Lateral Occipital Cortex, inferior division |
| 44 | -4.444 | -15.564 | 6.75 | 2.508 | 221 | JHA:<br>1% WM Fornix |
| 45 | 3.019 | -3.124 | 26.0 | 2.507 | 527 | JHA:<br>88% WM Callosal body, 3% WM Cingulum R<br><br>HOCSA:<br>9% Cingulate Gyrus, anterior division, 1% Cingulate Gyrus, posterior division |
| 45a | 0.531 | 4.339 | 28.75 | 2.347 | | JHA:<br>28% WM Callosal body<br><br>HOCSA:<br>54% Cingulate Gyrus, anterior division |
| 46 | 0.531 | 36.683 | 17.75 | 2.46 | 255 | HOCSA:<br>66% Cingulate Gyrus, anterior division, 10% Paracingulate Gyrus |
| 47 | 17.947 | 41.659 | -15.25 | 2.429 | 170 | HOCSA:<br>44% Frontal Pole, 2% Frontal Orbital Cortex |
| 48 | 27.899 | 26.731 | 1.25 | 2.408 | 289 | JHA:<br>2% WM Inferior occipito-frontal fascicle R<br><br>HOCSA:<br>10% Insular Cortex, 5% Frontal Orbital Cortex |
| 49 | 57.755 | -18.052 | 50.75 | 2.394 | 221 | JHA:<br>74% GM Primary somatosensory cortex BA1 R, 38% GM Primary somatosensory cortex BA2 R, 18% GM Inferior parietal lobule PFt R, 2% GM Primary somatosensory cortex BA3b R<br><br>HOCSA:<br>55% Postcentral Gyrus, 10% Supramarginal Gyrus, anterior division |

| | | | | | | |
|---|---|---|---|---|---|---|
| 50 | 7.995 | -77.764 | 39.75 | 2.386 | 221 | JHA:<br>30% GM Superior parietal lobule 7M R, 28% GM Superior parietal lobule 7P R<br><br>HOCSA:<br>37% Precuneous Cortex, 28% Cuneal Cortex, 1% Lateral Occipital Cortex, superior division |
| 51 | 17.947 | -50.396 | 23.25 | 2.38 | 187 | JHA:<br>49% WM Callosal body, 2% WM Optic radiation R<br><br>HOCSA:<br>12% Precuneous Cortex |

**Supplementary Table 5: Whole-brain clusters of the generalized Psychophysiological Interaction (gPPI) effect with SMA used as a seed region.** No clusters survived FDR correction in the whole-brain analysis using a voxel-level threshold of $p<0.01$. The table lists clusters uncorrected for multiple comparisons, with statistical significance threshold defined at a single voxel level of $z_{\alpha}=1.96$. Table displays MNI coordinates (X, Y, Z), statistical T values and atlas labels of peak voxels of the clusters. Atlas labels are based on the Juelich Histological Atlas (JHA) and Harvard-Oxford Cortical Structural Atlas (HOCSA). The labels were generated using FSL. Atlases are listed only if the labels were found for a given atlas. The letters after the cluster ID indicates its subcluster.

| Cluster ID | X | Y | Z | T Value | Cluster Size (voxels) | Atlas label |
|---|---|---|---|---|---|---|
| 1 | 32.875 | -40.444 | -9.75 | 4.459 | 1855 | JHA:<br>23% WM Optic radiation R, 7% GM Hippocampus cornu ammonis R, 6% WM Callosal body, 4% GM Hippocampus subiculum R, 4% GM Hippocampus dentate gyrus R<br><br>HOCSA:<br>38% Lingual Gyrus, 23% Temporal Occipital Fusiform Cortex, 12% Parahippocampal Gyrus, posterior division, 9% Temporal Fusiform Cortex, posterior division |
| 1a | 22.923 | -45.42 | -9.75 | 2.66 | | JHA:<br>1% GM Visual cortex V2 BA18 R<br><br>HOCSA:<br>68% Lingual Gyrus, 14% Temporal Occipital Fusiform Cortex, 1% Temporal Fusiform Cortex, posterior division |
| 1b | 27.899 | -55.372 | -7.0 | 2.057 | | JHA:<br>5% WM Optic radiation R, 1% GM Visual cortex V2 BA18 R<br><br>HOCSA:<br>37% Temporal Occipital Fusiform Cortex, 27% Lingual Gyrus, 6% Occipital Fusiform Gyrus |
| 2 | -49.228 | -5.612 | 45.25 | 4.247 | 5940 | JHA:<br>79% GM Premotor cortex BA6 L, 23% GM Primary motor cortex BA4a L, 8% WM Corticospinal tract L, 7% GM Primary somatosensory cortex BA1 L<br><br>HOCSA:<br>57% Precentral Gyrus, 3% Postcentral Gyrus |
| 2a | -49.228 | -5.612 | 53.5 | 3.663 | | JHA:<br>45% GM Premotor cortex BA6 L, 4% GM Primary motor cortex BA4a L, 2% GM Primary somatosensory cortex BA1 L<br><br>HOCSA:<br>55% Precentral Gyrus, 2% Middle Frontal Gyrus, 1% Postcentral Gyrus |
| 2b | -56.692 | -5.612 | 48.0 | 3.541 | | JHA: |

| | | | | | | |
|---|---|---|---|---|---|---|
| | | | | | | 39% GM Premotor cortex BA6 L, 9% GM Primary motor cortex BA4a L, 6% GM Primary somatosensory cortex BA1 L<br><br>HOCSA:<br>46% Precentral Gyrus, 2% Postcentral Gyrus |
| 2c | -61.668 | -13.076 | 26.0 | 3.208 | | JHA:<br>50% GM Secondary somatosensory cortex / Parietal operculum OP4 L, 26% GM Primary somatosensory cortex BA1 L, 22% GM Inferior parietal lobule PFop L, 16% GM Inferior parietal lobule PFt L, 10% GM Secondary somatosensory cortex / Parietal operculum OP1 L, 10% GM Primary somatosensory cortex BA3b L, 10% GM Primary somatosensory cortex BA2 L, 8% GM Primary somatosensory cortex BA3a L, 1% GM Broca's area BA44 L<br><br>HOCSA:<br>74% Postcentral Gyrus, 3% Supramarginal Gyrus, anterior division |
| 3 | 15.459 | 6.827 | 37.0 | 3.912 | 1293 | JHA:<br>29% WM Callosal body |
| 4 | -66.644 | -13.076 | -1.5 | 3.696 | 6655 | HOCSA:<br>45% Superior Temporal Gyrus, posterior division, 6% Middle Temporal Gyrus, anterior division, 5% Middle Temporal Gyrus, posterior division, 3% Superior Temporal Gyrus, anterior division, 1% Planum Temporale |
| 4a | -59.18 | 9.315 | 4.0 | 3.667 | | JHA:<br>38% GM Broca's area BA44 L, 10% GM Broca's area BA45 L<br><br>HOCSA:<br>20% Inferior Frontal Gyrus, pars opercularis, 17% Precentral Gyrus |
| 4b | -49.228 | 9.315 | 12.25 | 3.524 | | JHA:<br>35% GM Broca's area BA44 L, 2% GM Broca's area BA45 L<br><br>HOCSA:<br>40% Inferior Frontal Gyrus, pars opercularis, 10% Precentral Gyrus |
| 4c | -61.668 | -25.516 | 4.0 | 2.657 | | JHA:<br>10% GM Secondary somatosensory cortex / Parietal operculum OP1 L, 2% GM Primary auditory cortex TE1.0 L<br><br>HOCSA:<br>40% Superior Temporal Gyrus, posterior division, 14% Planum Temporale, 3% Middle Temporal Gyrus, posterior division |
| 5 | -31.812 | -18.052 | 23.25 | 3.608 | 1123 | JHA:<br>25% WM Superior longitudinal fascicle L, 14% GM Secondary somatosensory cortex / Parietal operculum OP3 L, 11% GM Secondary somatosensory cortex / Parietal operculum OP2 L, 9% WM Corticospinal tract L, 5% GM Insula Ig2 L<br><br>HOCSA:<br>1% Central Opercular Cortex |
| 5a | -26.836 | -10.588 | 31.5 | 3.231 | | JHA:<br>62% WM Corticospinal tract L, 3% WM Superior occipito-frontal fascicle L |
| 6 | -16.884 | -95.18 | -23.5 | 3.608 | 1174 | HOCSA:<br>10% Occipital Pole, 3% Occipital Fusiform Gyrus |
| 6a | -16.884 | -92.692 | -34.5 | 2.529 | | No label found |
| 6b | -31.812 | -92.692 | -20.75 | 2.515 | | JHA:<br>11% GM Visual cortex V3V L, 7% GM Visual cortex V4 L, 3% GM Visual cortex V2 BA18 L<br><br>HOCSA:<br>14% Lateral Occipital Cortex, inferior division, 11% Occipital Pole, 3% Occipital Fusiform Gyrus |
| 6c | -9.42 | -100.156 | -12.5 | 2.231 | | JHA:<br>71% GM Visual cortex V1 BA17 L, 38% WM Optic radiation L, 37% GM Visual cortex V2 BA18 L, 5% GM Visual cortex V3V L<br><br>HOCSA:<br>52% Occipital Pole, 2% Lateral Occipital Cortex, inferior division |
| 7 | 62.731 | -18.052 | -1.5 | 3.597 | 2996 | HOCSA: |

| | | | | | | |
|---|---|---|---|---|---|---|
| | | | | | | 49% Superior Temporal Gyrus, posterior division, 11% Middle Temporal Gyrus, posterior division, 2% Planum Temporale, 1% Superior Temporal Gyrus, anterior division |
| 7a | 70.195 | -13.076 | -1.5 | 3.519 | | HOCSA:<br>30% Superior Temporal Gyrus, posterior division, 3% Middle Temporal Gyrus, posterior division |
| 7b | 52.779 | -20.54 | -4.25 | 3.399 | | JHA:<br>1% GM Insula Id1 R<br><br>HOCSA:<br>53% Superior Temporal Gyrus, posterior division, 14% Middle Temporal Gyrus, posterior division |
| 7c | 70.195 | -8.1 | -1.5 | 2.996 | | HOCSA:<br>9% Superior Temporal Gyrus, posterior division, 2% Superior Temporal Gyrus, anterior division |
| 8 | -1.956 | 1.851 | 67.25 | 3.536 | 2212 | JHA:<br>74% GM Premotor cortex BA6 L<br><br>HOCSA:<br>72% Juxtapositional Lobule Cortex (formerly Supplementary Motor Cortex), 3% Superior Frontal Gyrus |
| 9 | -19.372 | -42.932 | -4.25 | 3.449 | 3455 | JHA:<br>22% GM Hippocampus subiculum L, 19% WM Callosal body, 12% WM Cingulum L, 2% WM Optic radiation L, 2% GM Hippocampus cornu ammonis L<br><br>HOCSA:<br>8% Lingual Gyrus, 7% Parahippocampal Gyrus, posterior division, 6% Cingulate Gyrus, posterior division |
| 9a | -21.86 | -35.468 | 1.25 | 3.205 | | JHA:<br>63% GM Hippocampus dentate gyrus L, 50% WM Fornix, 47% GM Hippocampus cornu ammonis L, 4% WM Callosal body, 1% GM Hippocampus subiculum L |
| 9b | -29.324 | -37.956 | -7.0 | 3.151 | | JHA:<br>78% GM Hippocampus cornu ammonis L, 28% GM Hippocampus dentate gyrus L, 15% WM Optic radiation L, 7% WM Callosal body, 4% WM Fornix<br><br>HOCSA:<br>18% Parahippocampal Gyrus, posterior division, 4% Lingual Gyrus, 2% Temporal Fusiform Cortex, posterior division, 1% Temporal Occipital Fusiform Cortex |
| 9c | -29.324 | -50.396 | -7.0 | 3.09 | | JHA:<br>4% WM Optic radiation L<br><br>HOCSA:<br>54% Temporal Occipital Fusiform Cortex, 17% Lingual Gyrus, 5% Temporal Fusiform Cortex, posterior division |
| 10 | 7.995 | -13.076 | -37.25 | 3.405 | 170 | No label found |
| 11 | 45.315 | 11.803 | -31.75 | 3.385 | 1719 | HOCSA:<br>60% Temporal Pole |
| 11a | 45.315 | 21.755 | -31.75 | 3.226 | | HOCSA:<br>53% Temporal Pole |
| 12 | -21.86 | -35.468 | -23.5 | 3.253 | 374 | HOCSA:<br>7% Parahippocampal Gyrus, posterior division, 6% Temporal Fusiform Cortex, posterior division |
| 13 | -4.444 | 46.635 | 56.25 | 3.168 | 783 | HOCSA:<br>1% Frontal Pole |
| 13a | 3.019 | 46.635 | 50.75 | 2.662 | | HOCSA:<br>20% Frontal Pole, 18% Superior Frontal Gyrus |
| 14 | 65.219 | -10.588 | 28.75 | 3.086 | 1872 | JHA:<br>28% GM Primary somatosensory cortex BA1 R, 24% GM Secondary somatosensory cortex / Parietal operculum OP4 R, 15% GM Primary somatosensory cortex BA3b R, 13% GM Primary somatosensory cortex BA2 R, 10% GM Inferior parietal lobule PFt R, 8% GM Secondary somatosensory cortex / Parietal operculum OP1 R, 7% GM Inferior parietal lobule PFop R, 4% GM Primary |

| | | | | | | |
|---|---|---|---|---|---|---|
| | | | | | | motor cortex BA4p R, 2% GM Inferior parietal lobule PF R, 1% GM Primary motor cortex BA4a R<br><br>HOCSA:<br>74% Postcentral Gyrus, 1% Precentral Gyrus |
| 14a | 55.267 | -8.1 | 53.5 | 2.963 | | JHA:<br>14% GM Premotor cortex BA6 R, 4% GM Primary somatosensory cortex BA1 R, 2% GM Primary somatosensory cortex BA3b R<br><br>HOCSA:<br>5% Precentral Gyrus, 4% Postcentral Gyrus |
| 14b | 57.755 | 1.851 | 45.25 | 2.124 | | JHA:<br>10% GM Premotor cortex BA6 R, 1% GM Primary somatosensory cortex BA3b R<br><br>HOCSA:<br>21% Precentral Gyrus, 1% Middle Frontal Gyrus |
| 15 | -24.348 | -57.86 | 1.25 | 3.083 | 1072 | JHA:<br>41% GM Visual cortex V2 BA18 L, 40% WM Optic radiation L, 39% GM Visual cortex V1 BA17 L, 22% WM Callosal body, 2% GM Visual cortex V4 L<br><br>HOCSA:<br>15% Lingual Gyrus, 7% Precuneous Cortex, 3% Intracalcarine Cortex, 1% Cingulate Gyrus, posterior division |
| 15a | -26.836 | -67.812 | -7.0 | 2.259 | | JHA:<br>31% GM Visual cortex V4 L<br><br>HOCSA:<br>46% Occipital Fusiform Gyrus, 9% Temporal Occipital Fusiform Cortex, 9% Lingual Gyrus |
| 16 | -26.836 | 21.755 | -26.25 | 3.049 | 323 | HOCSA:<br>38% Frontal Orbital Cortex, 1% Temporal Pole |
| 16a | -21.86 | 14.291 | -23.5 | 2.566 | | HOCSA:<br>78% Frontal Orbital Cortex |
| 17 | -36.788 | 16.779 | -4.25 | 3.042 | 1702 | HOCSA:<br>64% Insular Cortex, 2% Central Opercular Cortex, 1% Frontal Operculum Cortex |
| 17a | -39.276 | 24.243 | -7.0 | 2.992 | | HOCSA:<br>70% Frontal Orbital Cortex, 4% Frontal Operculum Cortex |
| 17b | -36.788 | 11.803 | 1.25 | 2.697 | | HOCSA:<br>58% Insular Cortex, 7% Frontal Operculum Cortex, 5% Central Opercular Cortex |
| 17c | -34.3 | 11.803 | 12.25 | 2.114 | | JHA:<br>8% GM Broca's area BA44 L<br><br>HOCSA:<br>44% Frontal Operculum Cortex, 14% Central Opercular Cortex, 8% Insular Cortex |
| 18 | -21.86 | 11.803 | 42.5 | 2.986 | 408 | HOCSA:<br>7% Superior Frontal Gyrus, 1% Middle Frontal Gyrus |
| 19 | -44.252 | -28.004 | 28.75 | 2.964 | 221 | JHA:<br>9% GM Secondary somatosensory cortex / Parietal operculum OP1 L, 5% GM Inferior parietal lobule PFop L, 2% GM Inferior parietal lobule PFt L<br><br>HOCSA:<br>3% Supramarginal Gyrus, anterior division, 2% Parietal Operculum Cortex, 2% Central Opercular Cortex |
| 20 | 0.531 | -57.86 | -23.5 | 2.872 | 1123 | No label found |
| 20a | 15.459 | -62.836 | -23.5 | 2.529 | | No label found |
| 20b | 0.531 | -57.86 | -34.5 | 2.28 | | No label found |
| 21 | 40.339 | 16.779 | -4.25 | 2.848 | 885 | JHA:<br>4% WM Inferior occipito-frontal fascicle R |

| | | | | | | |
|---|---|---|---|---|---|---|
| | | | | | | HOCSA:<br>73% Insular Cortex, 2% Frontal Operculum Cortex, 2% Frontal Orbital Cortex |
| 22 | -64.156 | 9.315 | 23.25 | 2.842 | 170 | JHA:<br>15% GM Broca's area BA44 L, 2% GM Premotor cortex BA6 L, 1% GM Broca's area BA45 L<br><br>HOCSA:<br>3% Precentral Gyrus |
| 23 | -9.42 | -10.588 | 37.0 | 2.842 | 697 | JHA:<br>26% WM Cingulum L, 21% WM Callosal body<br><br>HOCSA:<br>11% Cingulate Gyrus, anterior division, 1% Cingulate Gyrus, posterior division, 1% Juxtapositional Lobule Cortex (formerly Supplementary Motor Cortex) |
| 23a | -14.396 | -18.052 | 45.25 | 2.25 | | JHA:<br>22% WM Corticospinal tract L, 11% GM Premotor cortex BA6 L, 9% GM Superior parietal lobule 5Ci L<br><br>HOCSA:<br>8% Precentral Gyrus, 2% Cingulate Gyrus, posterior division, 1% Cingulate Gyrus, anterior division, 1% Juxtapositional Lobule Cortex (formerly Supplementary Motor Cortex) |
| 24 | -51.716 | -37.956 | -31.75 | 2.829 | 493 | HOCSA:<br>5% Inferior Temporal Gyrus, posterior division, 1% Inferior Temporal Gyrus, temporooccipital part |
| 24a | -51.716 | -37.956 | -42.75 | 2.304 | | No label found |
| 25 | -29.324 | 66.539 | 20.5 | 2.818 | 340 | HOCSA:<br>3% Frontal Pole |
| 25a | -34.3 | 59.075 | 26.0 | 2.694 | | HOCSA:<br>2% Frontal Pole |
| 26 | -54.204 | -77.764 | -4.25 | 2.815 | 340 | HOCSA:<br>25% Lateral Occipital Cortex, inferior division |
| 26a | -49.228 | -85.228 | -4.25 | 2.692 | | HOCSA:<br>13% Lateral Occipital Cortex, inferior division |
| 27 | 10.483 | 24.243 | 6.75 | 2.813 | 425 | JHA:<br>53% WM Callosal body |
| 28 | -26.836 | -15.564 | -18.0 | 2.802 | 817 | JHA:<br>88% GM Hippocampus cornu ammonis L, 58% GM Hippocampus dentate gyrus L, 31% GM Hippocampus subiculum L, 17% GM Amygdala_laterobasal group L, 14% GM Amygdala_superficial group L, 7% WM Fornix, 2% GM Hippocampus hippocampal-amygdaloid transition area L |
| 28a | -34.3 | -23.028 | -20.75 | 2.516 | | JHA:<br>68% GM Hippocampus cornu ammonis L, 26% GM Hippocampus dentate gyrus L, 14% WM Optic radiation L, 12% GM Hippocampus subiculum L, 1% WM Fornix<br><br>HOCSA:<br>21% Temporal Fusiform Cortex, posterior division, 17% Parahippocampal Gyrus, posterior division, 15% Parahippocampal Gyrus, anterior division |
| 29 | -1.956 | -35.468 | -59.25 | 2.784 | 238 | No label found |
| 30 | 0.531 | 44.147 | -1.5 | 2.747 | 272 | HOCSA:<br>44% Cingulate Gyrus, anterior division, 42% Paracingulate Gyrus, 2% Frontal Medial Cortex |
| 31 | -39.276 | 64.051 | -9.75 | 2.705 | 170 | HOCSA:<br>4% Frontal Pole |
| 32 | 25.411 | -85.228 | 31.5 | 2.698 | 306 | HOCSA:<br>48% Lateral Occipital Cortex, superior division, 24% Occipital Pole |

| | | | | | | |
|---|---|---|---|---|---|---|
| 33 | -29.324 | -23.028 | 56.25 | 2.69 | 187 | JHA:<br>61% WM Corticospinal tract L, 36% GM Primary motor cortex BA4p L, 31% GM Primary motor cortex BA4a L, 15% GM Premotor cortex BA6 L, 8% GM Primary somatosensory cortex BA3b L, 1% GM Primary somatosensory cortex BA3a L, 1% GM Primary somatosensory cortex BA1 L<br><br>HOCSA:<br>31% Precentral Gyrus, 17% Postcentral Gyrus |
| 34 | 32.875 | 11.803 | 53.5 | 2.684 | 187 | HOCSA:<br>34% Middle Frontal Gyrus, 5% Superior Frontal Gyrus, 1% Precentral Gyrus |
| 35 | 7.995 | -40.444 | -34.5 | 2.641 | 204 | No label found |
| 36 | -19.372 | 69.027 | 9.5 | 2.614 | 595 | HOCSA:<br>41% Frontal Pole |
| 36a | -29.324 | 66.539 | 4.0 | 2.534 | | HOCSA:<br>45% Frontal Pole |
| 37 | 57.755 | -35.468 | 56.25 | 2.614 | 663 | HOCSA:<br>16% Supramarginal Gyrus, posterior division, 1% Supramarginal Gyrus, anterior division |
| 38 | 3.019 | 14.291 | 12.25 | 2.508 | 510 | JHA:<br>12% WM Callosal body |
| 39 | -6.932 | -23.028 | 67.25 | 2.443 | 272 | JHA:<br>60% GM Premotor cortex BA6 L, 50% GM Primary motor cortex BA4a L, 39% WM Corticospinal tract L<br><br>HOCSA:<br>33% Precentral Gyrus |
| 39a | -11.908 | -15.564 | 67.25 | 2.401 | | JHA:<br>89% GM Premotor cortex BA6 L, 12% GM Primary motor cortex BA4a L, 11% WM Corticospinal tract L<br><br>HOCSA:<br>23% Precentral Gyrus, 5% Superior Frontal Gyrus, 1% Juxtapositional Lobule Cortex (formerly Supplementary Motor Cortex) |
| 40 | 37.851 | 34.195 | -23.5 | 2.416 | 204 | HOCSA:<br>4% Frontal Orbital Cortex, 2% Frontal Pole |
| 41 | -34.3 | -3.124 | -1.5 | 2.41 | 204 | JHA:<br>7% WM Inferior occipito-frontal fascicle L<br><br>HOCSA:<br>1% Insular Cortex |
| 42 | -11.908 | -37.956 | 70.0 | 2.361 | 204 | JHA:<br>38% GM Primary motor cortex BA4a L, 26% WM Corticospinal tract L, 22% GM Primary somatosensory cortex BA3b L, 16% GM Primary motor cortex BA4p L, 14% GM Superior parietal lobule 5L L, 13% GM Premotor cortex BA6 L, 10% GM Primary somatosensory cortex BA3a L, 9% GM Superior parietal lobule 5M L, 9% GM Primary somatosensory cortex BA1 L, 8% GM Primary somatosensory cortex BA2 L<br><br>HOCSA:<br>43% Postcentral Gyrus, 9% Precentral Gyrus, 2% Superior Parietal Lobule |
| 43 | -29.324 | -85.228 | 31.5 | 2.28 | 255 | JHA:<br>30% GM Inferior parietal lobule PGp L<br><br>HOCSA:<br>64% Lateral Occipital Cortex, superior division, 9% Occipital Pole |
